# Enzymes can activate and mobilize the cytoplasmic environment across scales

**DOI:** 10.1101/2025.01.28.635259

**Authors:** Mirco Dindo, Jakob Metson, Weitong Ren, Michalis Chatzittofi, Kiyoshi Yagi, Yuji Sugita, Ramin Golestanian, Paola Laurino

## Abstract

Biomolecular condensates have so far been studied in terms of their structural, compositional, and functional properties. However, condensate enzymatic activity —a key aspect of cellular metabolism— remains unexplored due to the complexity of the system. In this study, using a combination of experimental, computational and theoretical techniques, we have discovered that the non-equilibrium activity which originates from catalytic reactions couples with the environment through various feedback mechanisms across five orders of magnitude of length scales. We observe that condensed enzymes catalyse more rapidly in the presence of crowding proteins and show that the increased enzymatic activity within these droplets stems from the emergence of lower-energy protein conformations induced by the highly crowded environment. Despite the crowding in the environment of the droplet, which might suggest an effective increase in its overall viscosity, we find that it becomes more agile, as evidenced by the observation of enhanced diffusion and macroscopic flow, due to the enzymatic activity. These findings shed new light on the dynamic interplay between enzymatic activity, composition and crowding in condensates, and their roles on the mobility and accessibility of various functional units in these environments, offering a novel perspective on liquidliquid phase separation in metabolically active conditions.

## MAIN TEXT

The interior of a living cell is a highly structured and crowded milieu packed with proteins, nucleic acids, and smaller molecules [1–5]. Moreover, it hosts a plethora of metabolic activities – through biochemical reactions – which drive the system away from thermodynamic equilibrium and maintain it in its functioning homeostatic state. While these structural and metabolic aspects that underlie the biological function of the cell have been subject to extensive investigations, the relationship between them, which will be governed by the physical properties of the non-equilibrium active environment [6, 7], has so far remained largely unexplored.

Particularly intriguing is the potential to have synergies between the structure of the cell, its metabolic activity, and its ability to function. For example, it has been reported that the bacterial cytosol exhibits liquid-like behaviour in metabolically active states, while it transitions to having glass-like behaviour in dormant states [8]. The physical properties of the medium can be influenced, e.g. by dynamic conformational changes in the enzymes while undergoing the catalytic cycles [9, 10], as well chemical gradients that are produced and maintained in the course of the catalytic activity [11, 12], which can control the non-equilibrium interactions between proteins [13], and consequently, the glassiness of metabolically active protein condensates [14]. These non-equilibrium interactions can give rise to novel structure formation resulting from catalytic activity [15], and may have played a key role in the spontaneous formation of metabolic cycles at the origins of life [16, 17]. Moreover, the cell cycle dynamics is known to have mechanisms in place that enable the formation of enzyme-rich condensates and metabolic autoregulation [18, 19], possibly taking advantage of spatial organization that can arise from catalysis-induced non-equilibrium activity in protein condensates [20].

To elucidate such synergies it is imperative to combine mechanistic descriptions of the essential constituents of the system across scales. Enzymes are known to adopt multiple structural states in the form of conformational ensembles that are crucial for their catalytic activity, although many functionally relevant excited states of enzymes remain inaccessible and sparsely populated [21–25]. Crowding can strongly impact enzyme conformation and function via a variety of mechanisms[26], such as excluded volume effects that can influence the mobility of macromolecules [1, 27] and promote oligomerization [28–30]. Moreover, crowding can also compromise the stability of native enzymes via non-specific interactions [31–33], while restricting the encounters between macromolecules [34–37], which can affect the catalytic efficiency of diffusion-limited reactions [38, 39]. It is thus unclear how enzyme condensates will function under the crowded conditions and whether such effects can be used for regulation of their metabolic activity.

Recent studies discuss the possibility that catalytic reactions can influence enzyme diffusion [40–42]. Early experimental reports of enhanced enzyme diffusion, however, have been inconclusive [43, 44], as other mechanisms that are not related to enhanced diffusion, such as flurophore detachment [45] or aggregation effects [46, 47], can explain the reported observations. Indeed, alternative experiments on single enzymes (not using florescence-based techniques) [48] and even experiments on enzymes in crowded biological environments [49] have shown normal diffusion that agrees well with the Stokes-Einstein relation. One important aspect in analysing such experiments concerns the question of how to choose the correct base-line value for the diffusion coefficient in mixtures, as has been revealed over the course of the controversy surrounding the NMR measurements in connection with enhanced diffusion question [50–54]. Therefore one should draw conclusions with caution and consider any possible experimental artefacts. Notwithstanding these considerations, enhanced diffusion of enzymes due to catalytic activity is a remarkable theoretical possibility, and it would be pertinent to investigate whether it can be realized in enzyme condensates, and potentially exploited towards biological function.

Here, we study liquid-liquid phase-separation (LLPS) in metabolically active membraneless protein droplets, and uncover physicochemical feedback mechanisms that activate and mobilize the environment, encompassing a multitude of length scales from 1 Å to 10 µm (see Fig. 1). We find that the overall metabolic activity of the condensate – containing Lactate dehydrogenase (LDH) – exhibits strong and non-monotonic dependency on the degree of crowding. Our computational studies on LDH rationalize the observation, by revealing that different conformations of the enzyme can be stabilized under dilute and protein-crowded conditions. We find evidence suggesting that the enzymatic activity leads to the emergence of enhanced diffusion of particles in the condensate as well as a global flow, reminiscent of cytoplasmic streaming. Therefore, our study mechanistically demonstrates how enzymes can exploit detailed changes in their conformation to significantly influence their surrounding environment.

**FIG. 1.**
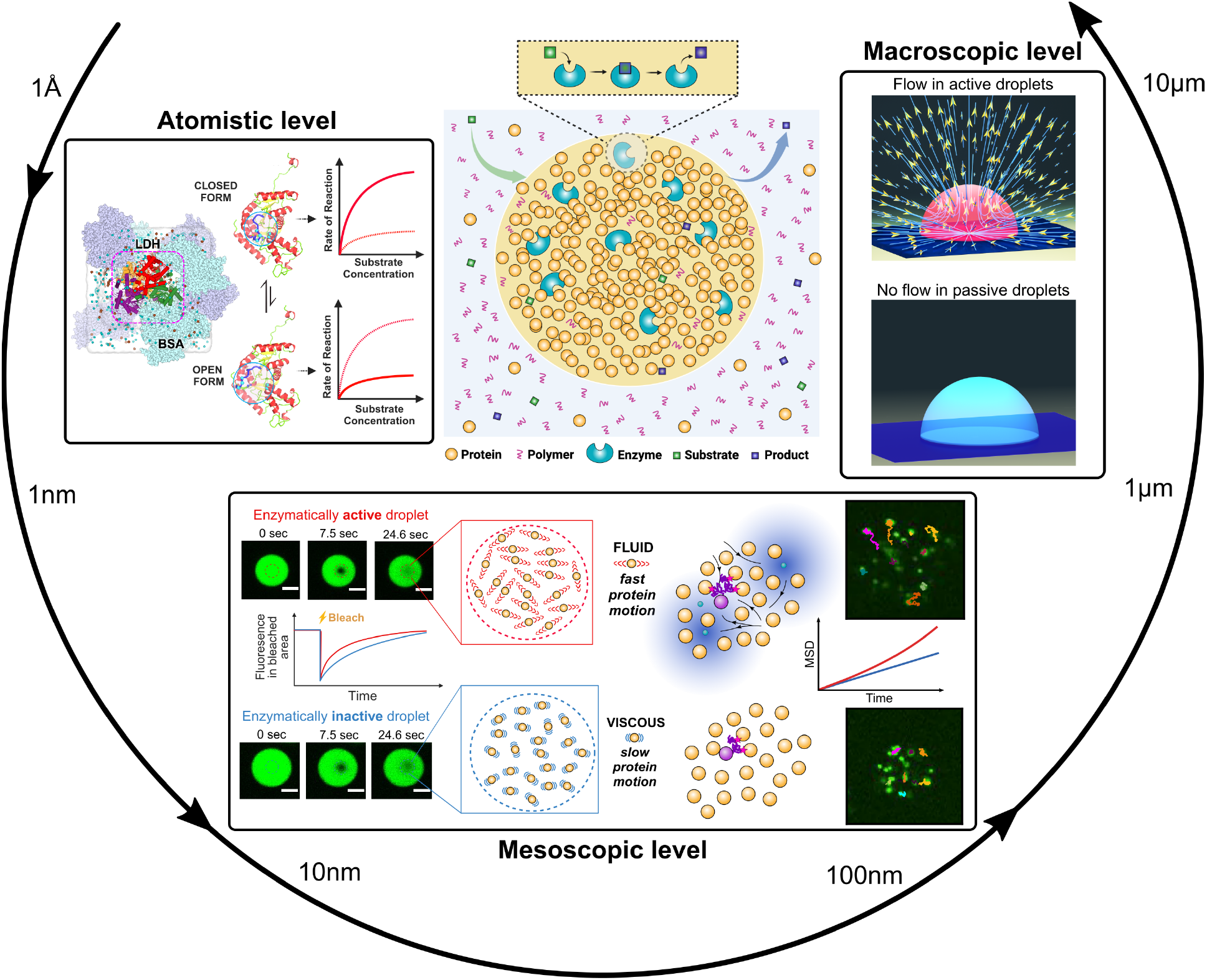
Interplay between enzymes and the crowded cytoplasmic environment across the scales. Crowded protein droplets allow the partitioning of enzymes to reach high enzymatic activity compared to dilute conditions. Computational dissection of enzyme conformations reveals the atomistic changes in the presence of crowders. Fluorescently tagged crowder proteins and nanoparticle tracking experiments indicate that there is faster movement in the crowded droplet environment in the presence of enzymatic activity. Large-scale flows in the droplets emerge due to chemical gradients produced by enzymes.

### Enzymatic activity increases mobility within the highly crowded protein droplets

To explore the impact of enzymatic activity on mobility within the droplets, we employed two methods: fluorescence recovery after photobleaching (FRAP) and particle tracking. In the FRAP experiments, we have used droplets containing different bovine serum albumin (BSA) concentrations (Fig. S1). Importantly, a small amount of BSA (5-10%) was labeled with Alexa Fluor 488, which served as the fluorescent marker. By utilizing the protein-crowded droplet system, we could precisely control and achieve high levels of enzymatic activity by adjusting both enzyme and substrate concentrations. We have selected L-lactate dehydrogenase (LDH) and yeast alcohol dehydrogenase (ADH), which are extensively studied oxidoreductases of broad biotechnological and metabolic interest [55, 56]. Previous work had indicated LDH’s partitioning and its activity in the droplets [11], which was confirmed for LDH (Fig. S2**a**) and further for ADH in droplets with different BSA concentrations in this study (Fig. S2**b**). Following photobleaching of the droplets, the fluorescence fully recovered in around 100 seconds, confirming the liquid-like behaviour of the droplets. However, due to the high protein crowding, the viscosity is approximately 2000-fold higher than that of water [11], akin to cytoplasmic measurements [57, 58]. Remarkably, a significant disparity in fluorescence recovery rate was observed between active and inactive droplets. LDH-active droplets, in the presence of 0.5 mM pyruvate, exhibited a 50% fluorescence recovery within approximately 8 seconds (7.62 ± 0.55 s), whereas inactive droplets (containing no pyruvate as substrate) achieved the same recovery after 14 seconds (13.23 ± 0.41 s), which can be seen in Fig. 2**a**. This discrepancy underscores that enzymatically active droplets recover fluorescence faster than inactive ones. To validate this observation, FRAP experiments were conducted by varying the LDH substrate concentration (0.25–1mM). As shown in Fig. 2**b**, increasing substrate concentration was found to be correlated with decreased FRAP recovery half-life values, indicating faster movement of BSA molecules in the presence of heightened enzymatic activity. Notably, when using droplets without any active enzyme but with increasing substrate or product concentration (0–1 mM substrate and 0–1 mM product), no variation in half-life values was observed Fig. E1, ruling out a passive chemical gradient influence and directly linking the changes to high enzymatic activity. Also, concentrations of substrates and cofactors have been quantified in both the continuous and droplets phases Fig. E3 and no differences in concentrations have been measured between the two phases.

**FIG. 2.**
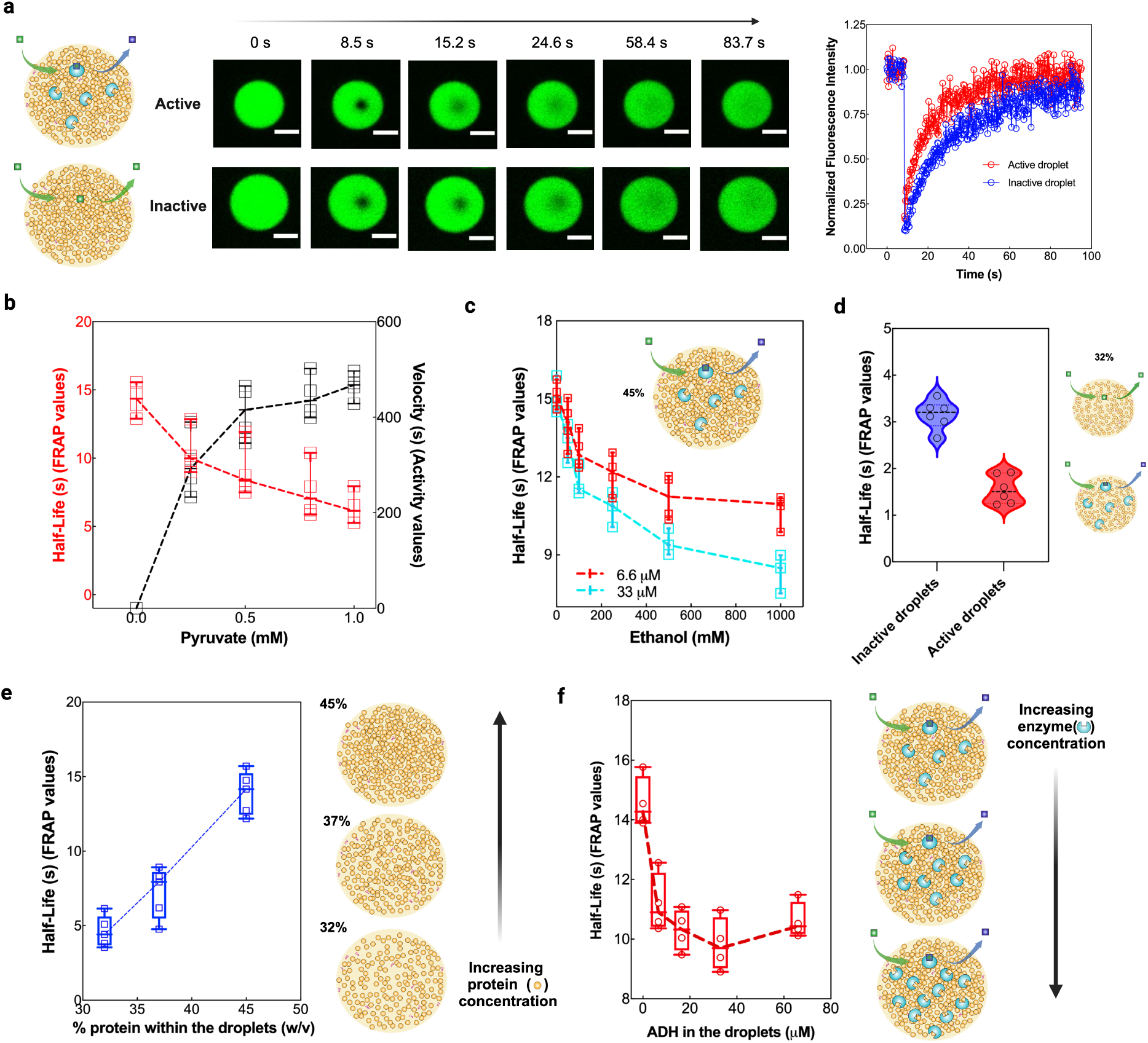
Enzymatic activity reveals changes of the crowded environment. **a**, Screenshot of the fluorescence recovery after photobleaching (FRAP) experiments on enzymatically active (red, in presence of active LDH) and inactive (blue, in absence of LDH) droplets using 0.5 mM pyruvate and 10 mM NADH. LDH concentration within the droplets was 3.3 µM. **b**, Trend of the half-life values expressed in seconds (red box and whiskers) and reaction rate (black box and whiskers) of LDH using 3.3 µM as a final concentration inside the droplets, as a function of substrate concentration (pyruvate concentration from 0 to 1.0 mM) The droplets dilution before the experiments has been the same for both the experiments (activity and FRAP). **c**, Trend of the BSA half-life values (expressed in seconds) using active ADH at two different enzyme concentrations within the droplets (red 6.6 µM and cyan 33 µM) as a function of substrate concentration (ethanol, 0-1 M). **d**, BSA half-life values (expressed in seconds) obtained using droplets containing 32% BSA w/v (around 4-5 mM protein concentration), in absence (blue box and whiskers, inactive droplets) and in presence (red box and whiskers, active droplets) of 500 mM ethanol as substrate using droplets with partitioned 6.6 µM ADH in presence of 10 mM NAD+. **e**,Trend of the half-life (expressed as seconds) using inactive droplets with different BSA content (32%, 37% and 45% w/v). No enzymatic activity was present in these experiments. **f**, Trend of the BSA half-life values using active ADH at different concentrations within the droplets (from 6.6 µM to 66 µM) in presence of a fixed ethanol 500 mM and in presence of 10 mM NAD+. All the experiments have been performed at least in triplicate (n = 3) and are reported as mean ± standard deviation. Data analysis was performed using GraphPad Prism 10.4.1.

To further elucidate the effect of enzymatic activity and different crowding concentrations on mobility in the droplet phase, FRAP experiments were performed with ADH at two concentrations (6.6 µM and 33 µM). Active ADH induced faster fluorescence recovery in droplets, with the higher ADH concentration leading to even faster recovery rates (Fig. 2**c**). This suggests that increased enzymatic activity prompts a “fluidification” effect relative to BSA within droplets, shifting toward a more liquidlike state. Moreover, experiments with droplets containing 32% w/v BSA and 500 mM ethanol added to the ADH showed decreased half-life values compared to inactive ADH-containing droplets (Fig. 2**d**), corroborating earlier observations with LDH in more crowded droplets (45% w/v BSA) (Fig. 2**b**). Further experiments using droplets containing varying BSA concentrations revealed that increasing protein content led to gradual increases in half-life values (about 3- and 4-folds), in line with expectations due to higher protein concentration and crowding (Fig. 2**e**). Interestingly, FRAP experiments with increasing enzymatic concentrations (ADH up to 66 µM) revealed that the fluidification effect mediated by enzymatic activity was detectable only up to 33 µM ADH (Fig. 2**e**). Above this concentration, no changes in halflife values were observed. The effect of metabolic density on the droplet’s physical state was most pronounced with 6.6 µM ADH, decreasing the half-life by 2.7 seconds (from 13.8 ± 0.2 to 11.2 ± 0.5 seconds). In summary, even low enzymatic activity within the droplets triggers fluidification effects, suggesting a strong impact on the physical state mediated by enzymatic activity.

We extract diffusion coefficients for the BSA proteins from the FRAP experiments by constructing a theoretical model FRAP system, as shown in Fig. 3**a** and Fig. 3**b**. As expected from the decreased FRAP half-life in active droplets, we find a higher diffusion in the active droplets. In particular, in the inactive droplets we find *D*_BSA_ = (9.3 ± 0.3) × 10^*−*3^ µm^2^ s^*−*1^ and in active droplets we find *D*_BSA_ = (2.3 ± 0.1) × 10^*−*2^ µm^2^ s^*−*1^. These results are summarised in Fig. 3**c**.

**FIG. 3.**
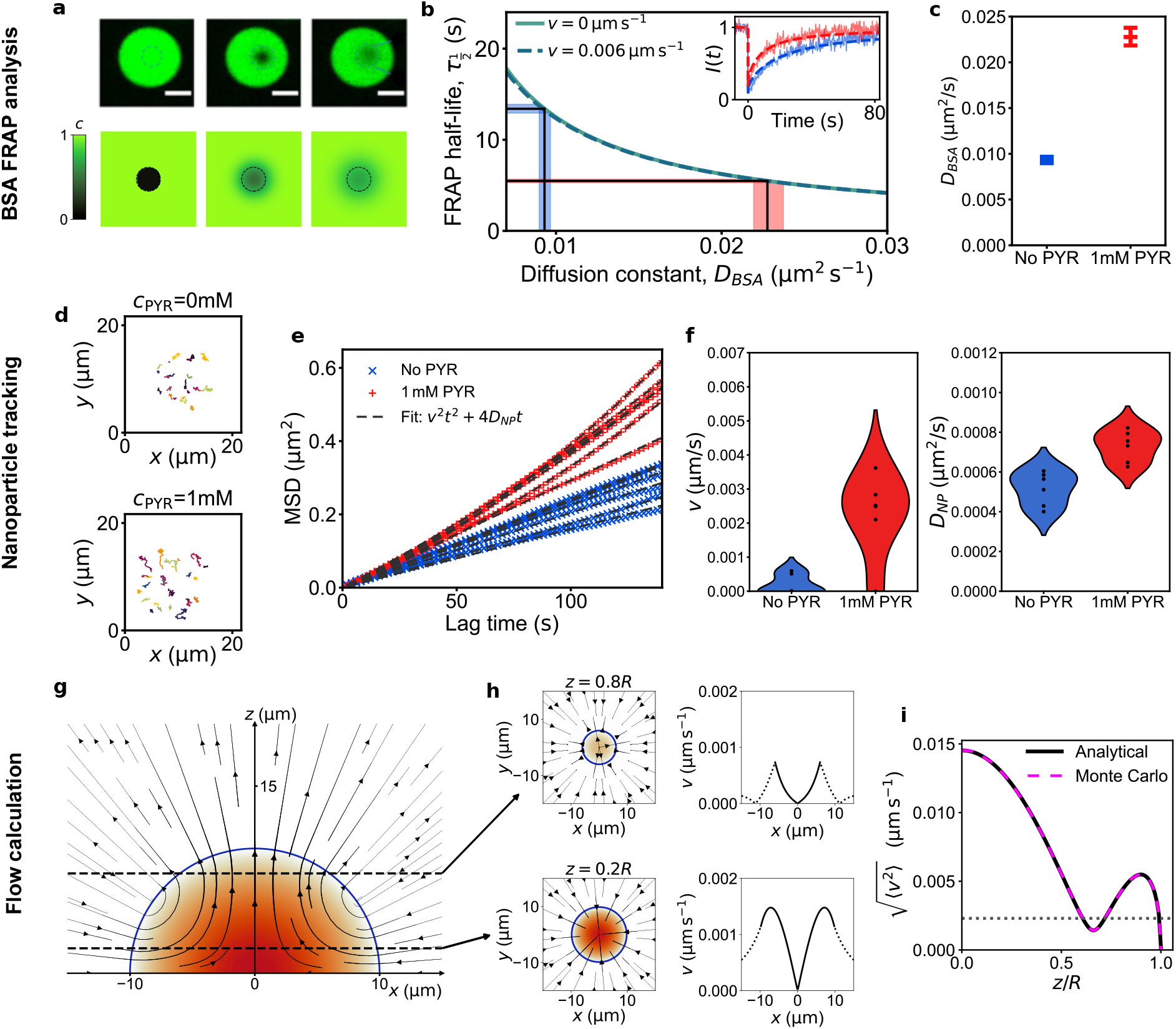
Effect of enzymatic activity on the droplet environment. **a**, Top: Microscope images of fluorescence in a real droplet following a bleach. Bottom: Fluorescence in the simulated system following a bleach. **b**, FRAP recovery half-life as a function of the diffusion coefficient. As shown by the dashed curve, the flow velocities measured in tracking have a negligible effect at this scale. Shaded bars show the inversion method used to obtain diffusion coefficients given the experimentally measured half-lives. Inset: Simulated intensity recovery curves plotted over experimental curves. **c**, Diffusion coefficients for the BSA crowder proteins in inactive and in active droplets. **d**, Nanoparticle trajectories measured in an inactive droplet (top) and in an active droplet (bottom). **e**, Mean-squared displacements of nanoparticle tracers measured in six inactive droplets (blue) and six active droplets (red). **f**, Flow velocities *v* and diffusion coefficients *D* for the nanoparticles in active and inactive droplets extracted from the mean-squared displacement measurements. **g**, Analytically calculated flow profile within and around a droplet due to enzymatic activity (Eqs. (2) and (3)), with the density profile shown as a heatmap (Eq. (1)). **h**, Horizontal flow profiles and horizontal flow velocity *v* shown at different heights (see SI). **i**, Variation of the root-mean-squared horizontal flow velocity within the droplet as a function of height (see SI). The analytical result has been verified using a Monte Carlo calculation. The dotted horizontal line shows the average value measured from experiments. This calculation shows a good agreement with our experiments without using any adjustable parameters.

Tracking the motion of nanoparticle tracers in droplets provides an alternative, independent method to measure mobility in the crowded environment. As can be seen in Fig. 3**e**, the mean-squared displacement (MSD) of nanoparticles is higher in active droplets. From the MSD of the tracers we can quantitatively extract the drift and diffusion parameters. The extracted parameters are shown in Fig. 3**f**. The diffusion coefficients we measure are of the same order as those measured previously in similar droplets [11]. Furthermore, the average diffusion constant of the nanoparticles is significantly higher in the active droplets. For inactive droplets we obtain *D*_NP_ = (5.2 ± 0.8) × 10^*−*4^ µm^2^ s^*−*1^, whilst in active droplets we measure *D*_NP_ = (7.3 ± 0.7) × 10^*−*4^ µm^2^ s^*−*1^. There is no measurable drift velocity in the inactive droplets, and in the active LDH droplets we find a drift velocity of *v* = (2.3 ± 1.1) × 10^*−*3^ µm s^*−*1^. We have checked the robustness of our extract parameters with respect to changes of the thresholds in our data analysis, as shown in Extended Data Fig. E2.

A provided theoretical calculation which is in good agreement with the tracking analysis, serves as a simple non-equilibrium mechanism for explaining the drift flow velocity inside the active droplet as indicated in Fig. 3**g**-**i**. Our calculation of these flow profiles (see Methods and SI) given in Eqs. (2) and (3) as well as the overall scale of the velocity given in Eq. (4) agree very well with our observations without the use of any adjustable parameters. Our theory predicts that the enhanced agility in metabolically active droplets is due to the interplay between chemical gradients – as created by the combined effect of enzyme condensation and enzyme metabolic activity – and the interaction with the solid surface that the condensate is adjacent to – as a confining boundary – both of which are universally generic ingredients that will be present in any experimental condition. In simple terms, this model suggests that the interaction with the surface creates hydrodynamic flows that engage with the movement of any constituent that is dissolved in the “synthetic cytosol”, as if the condensate is a mixing chamber.

### Protein crowding influences single enzyme kinetic parameters

The enzymatic parameters were measured to investigate the effect of macromolecular crowding in the droplets (I). LDH catalytic efficiency within droplets surpassed that of diluted conditions (*k*_cat_*/K*_M_ = 1253 ± 307 s^*−*1^ mM^*−*1^) from 1-fold (*k*_cat_*/K*_M_ = 2700 ± 729 s^*−*1^mM^*−*1^ using highly crowded droplets with BSA 45% w/v) [11] to 4-fold (*k*_cat_*/K*_M_ = 5433 ± 1412 s^*−*1^ mM^*−*1^ using less crowded droplets with BSA 32% w/v), driven by increased turnover number values *k*_cat_ (from 320 ± 39 s^*−*1^ in buffer to 815 ± 65 s^*−*1^ in droplets with 32% w/v BSA) and decreased *K*_M_ values (from 0.26 ± 0.06 mM in buffer to 0.15 ± 0.04 mM using droplets with BSA 32% w/v) (I). Similarly, ADH exhibited 3-fold enhanced catalytic efficiency in droplets shifting the *k*_cat_*/K*_M_ from 12 ± 3 s^*−*1^ mM^*−*1^) to 35 ± 6 s^*−*1^ mM^*−*1^ using droplets with 32% w/v, with increased *k*_cat_ values (from 450 ± 33 s^*−*1^ in buffer to 904 ± 61 s^*−*1^ in 32% BSA droplets) and slightly decreased *K*_M_ values (from 34 ± 3 mM in buffer to 26 ± 4 mM in droplets). These findings suggest an overall enhancement of enzyme catalytic efficiency in droplets compared to dilute conditions Fig. S3**a** [59]. Importantly, in the absence of LDH and ADH, substrate and product concentrations were equal across the droplet and continuous phases as reported in the Fig. E3, indicating unhindered diffusion between the phases and linking changes in kinetic parameters to the presence of enzyme. Trends of *k*_cat_ and *K*_M_ values against BSA concentration in the droplets (Fig. S3**b**) showed that decreasing protein concentration inside the droplets (from 45% to 32% w/v BSA) led to increased *k*_cat_ values for both enzymes analyzed (between 1- and 2-folds). Additionally, the *K*_M_ value of LDH decreased (1-fold), suggesting increased substrate affinity, while the *K*_M_ value of ADH for ethanol remained mostly unchanged (I).

**TABLE 1.** Kinetic parameters values of the enzyme studied in buffer and in the phase separated crowded droplets containing different BSA concentrations. Values of *k*_cat_, *K*_M_, and *k*_*cat*_*/K*_*m*_ have been measured in triplicate and by using different concentration of pyruvate (0-1mM) for LDH and ethanol (0-1M) for ADH. Concentration of free enzyme and within the droplets was the same for all the measurements. LDH concentration withing the droplets was 3.3 µM and for ADH was 6.6 µM. The kinetic parameters have been measured as reported in the Material and Methods section.

| LDH | $k_{\text{cat}}$ ($\text{s}^{-1}$) | $K_{\text{M}}$ (mM) | $k_{\text{cat}}/K_{\text{M}}$ $\text{s}^{-1} \text{mM}^{-1}$ |
| --- | --- | --- | --- |
| Buffer | $320 \pm 39$ | $0.26 \pm 0.06$ | $1253 \pm 307$ |
| 45% | $540 \pm 61$ | $0.20 \pm 0.05$ | $2700 \pm 729$ |
| 37% | $777 \pm 69$ | $0.16 \pm 0.04$ | $4856 \pm 1262$ |
| 32% | $815 \pm 65$ | $0.15 \pm 0.04$ | $5433 \pm 1412$ |
| ADH | $k_{\text{cat}}$ ($\text{s}^{-1}$) | $K_{\text{M}}$ (mM) | $k_{\text{cat}}/K_{\text{M}}$ $\text{s}^{-1} \text{mM}^{-1}$ |
| Buffer | $450 \pm 33$ | $34 \pm 3$ | $12 \pm 3$ |
| 45% | $716 \pm 45$ | $25 \pm 4$ | $26 \pm 5$ |
| 37% | $830 \pm 55$ | $26 \pm 5$ | $32 \pm 6$ |
| 32% | $904 \pm 61$ | $26 \pm 4$ | $35 \pm 6$ |

The overall effect of compartmentalization and crowding within droplets on enzyme catalytic efficiency is depicted in Fig. S3**c**. Compartmentalization within liquid droplets with high protein crowding greatly enhanced overall catalytic efficiency compared to dilute conditions Fig. S3**c**. However, further increasing the protein concentration within the droplets lead to a decrease in catalytic efficiency (3-folds). The enhancement of catalytic activity mediated by crowding is typically attributed to the excluded volume effect, related to nonspecific steric and repulsive interactions. However, the observed non-monotonic trend indicates that a certain degree of freedom is needed for the enzyme to maintain its catalytic efficiency and dynamics. Thus, it is not solely the excluded volume contributing to increased catalytic efficiency, but also thermodynamic and conformational factors related to the enzyme and the environment.

### Atomistic mechanisms for the crowding effects on enzyme activities

All-atom molecular dynamics (MD) simulations of the apo state of LDH in dilute conditions revealed the activesite loop and the contacting helix are mostly separated (r(loop-helix)=17 Å) (Fig. S4). The substrate binding in the holo state of LDH induces a meta-stable half-open form (r(loop-helix)=12 Å) in addition to the closed form. In crowded (8bsa) conditions, the closed form (r(loop-helix)=7.5 Å) appears in both apo and holo states, while the stability of the closed form in holo state is much greater than that in apo state. Thus, the crowding conditions significantly stabilize the closed form of the activesite conformations in LDH. In the open form of X-ray crystal structure of the LDH ternary complex (chain A in 3H3F), oxamate, a ligand mimicking the effect of the pyruvate binding, interacts with Asn137, Arg168, His192, and Thr247. Arg105 additionally stabilizes the oxamate binding by adding two more hydrogen bonds in the closed form (chain B in 3H3F)), suggesting the functional roles of Arg105. Indeed, Arg105 and pyruvate were frequently observed within the hydrogen-bonded distance (3.5 Å) in MD simulations of the holo state in crowded (8bsa) conditions (Fig. S5). The hydrogen bonds seem to impose additional structural restraints not only to pyruvate but also to the proper reactive poses of His192 and NADH, which are proton and hydride donors to the pyruvate in the LDH catalytic reactions (Fig. E4**b**).

Hybrid quantum mechanics/molecular mechanics (QM/MM) calculations of hydride transfer (HT) and proton transfer (PT) reactions in the active site of LDH, as illustrated in Fig. 4**c**, were performed using the open and closed forms of LDH. The computed potential of mean forces (PMFs) exhibits only one free-energy barrier, suggesting a concerted mechanism for the enzyme reaction [60] (Fig. 4**d**). Interestingly, the closed form exhibits a lower barrier height and a more stable product state. 2D-PMF of the closed form in terms of HT and PT coordinates (r(HT) and r(PT), respectively) reveals that the HT reaction precedes the PT reaction along the minimum energy path (Fig. 4**e**, see Fig. S6 for the open form). The bond alternation points of HT and PT reactions (r(HT) = 0 and r(PT) = 0; (Fig. S7) and Fig. E5) correspond to s ∼ 0.5 and 0.8, respectively, where the central barrier and a small shoulder are observed in the 1D-PMF (Fig. 4**d**). Therefore, the hydrogen bonds between Arg105 and pyruvate primarily contribute to reducing the barrier height of HT reaction, thereby enhancing the catalytic activity.

**FIG. 4.**
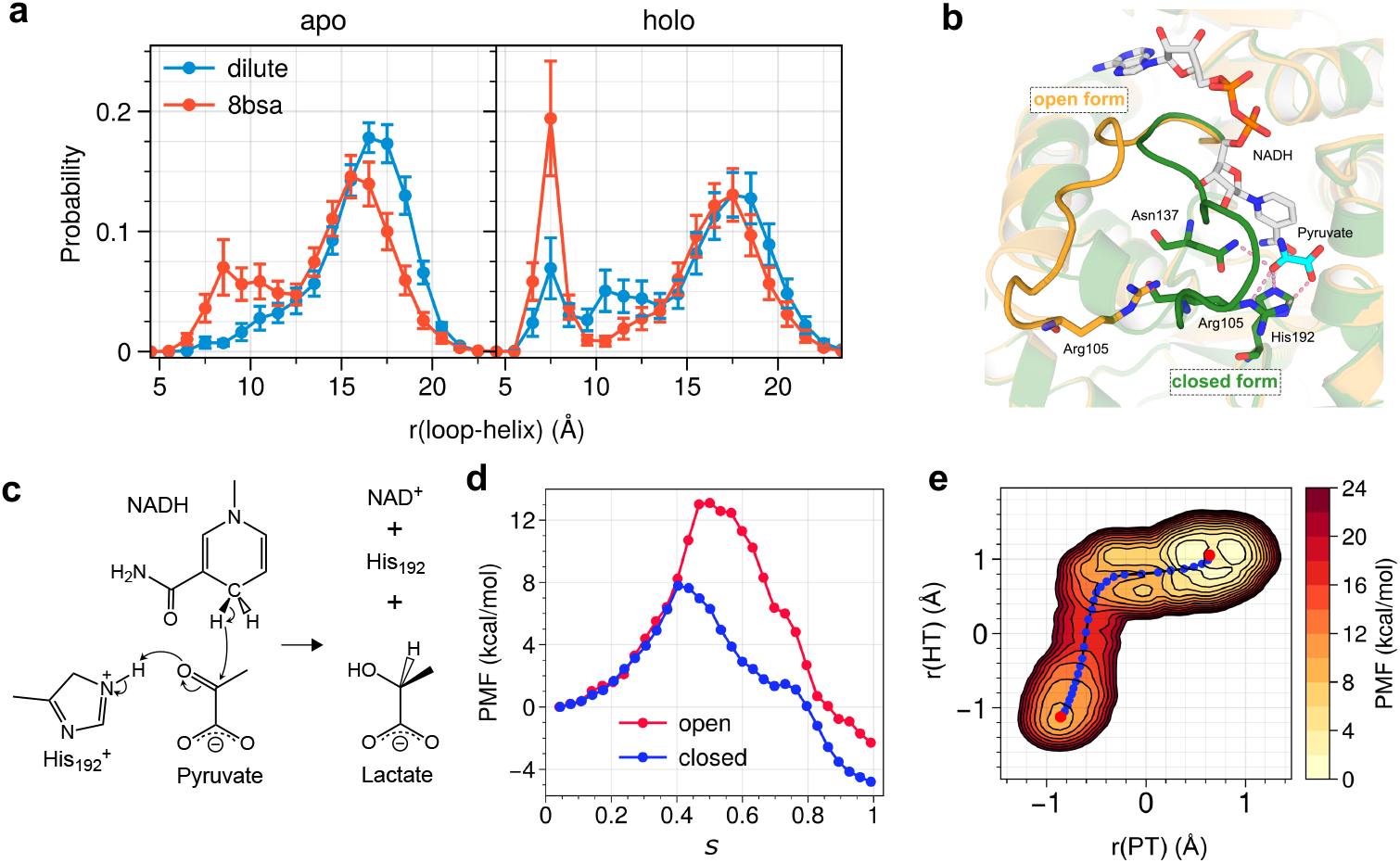
Enzymatic reaction mechanisms in dilute and crowded environment. **a**, Histograms of loop-helix distance in the apo (left) and holo (right) states in dilute and crowded (8bsa) conditions. The loop-helix distance is defined as the distance between the COM of C*α* atoms of residues 101-103 in the active site loop and that of Ca atoms of residues 123-124 in its contacting helix. **b**, 3D structures of the binding pocket with the loop in open (orange) and closed (green) forms, the ligand (pyruvate), NADNH, and sidechains of relevant amino acid residues (Arg105, Asn137, His 192). **c**, A hydride transfer (HT) from NADH and a proton transfer (PT) from His192 to pyruvate yields lactate. **d**, 1D-PMF along a reaction coordinate in the open and closed states. **e**, 2D-PMF in terms of PT and HT coordinates in the closed state.

The QM/MM simulations suggest the functional importance of Arg105 in the closed form of LDH, which is stabilized in crowded (bsa8) conditions, as shown in all-atom MD simulations. The crowding effects on enzyme LDH result not from static but highly dynamic interactions between LDH and BSA crowders (Fig. S8). We note that in crowded (bsa8) conditions, diffusions of substrates (pyruvate and NADH) are slowed down as the volume fraction of BSA increases (Fig. S9). Pyruvate is a small molecule with one negative charge, while NADH is a larger molecule with two negative charges. Both are observed to rapidly absorb on BSA surfaces, forming multiple hydrogen bonds with BSA molecules (see Fig. S10). Macromolecular crowding effects on enzyme catalysis were discussed in terms of the transition-state stabilization (positive effect) and the slow-down of substrate diffusion (negative effect) [1, 29, 61]. Here, we could observe both effects and explain the experimental observations of LDH enzyme catalysis as a function of BSA content in the droplets (Fig. S3**c**).

We thus find that the transition state of the closed form of LDH, which is preferred in a crowded environment, shows a lower activation energy than that of the open form, explaining the higher catalytic activity.

## Discussion

To understand how macromolecular crowding influences enzymatic parameters, investigations were conducted in highly crowded environments using a liquid-liquid phase separated system. The system allows for the isolation and study of specific elements, such as protein crowding and enzymatic activity, within minimal chemical systems, reducing biological complexity while preserving physiological relevance. Through the manipulation of single components, it was demonstrated that enzymatic activity, mediated by LDH and ADH, facilitates the movement of BSA proteins and nanoparticle tracers in the highly crowded protein droplets. This effect was observed to be independent of the presence of small molecules like substrate, cofactors, or reaction products, suggesting a direct link to enzyme dynamics during catalysis.

Based on the FRAP and tracking results, we propose the following picture that can explain the low diffusion coefficient in the droplets and the relative enhancement in the active droplets. Considering the dense complex environment of the droplets, it is expected that diffusion will be hindered by local interactions between the particles in the crowded environment. To incorporate this physical picture, we propose the following simple model. Particles can be in one of two states: either able to diffuse or significantly slowed down (or immobilised) due to interactions with other particles in the droplets. This model is illustrated in Fig. 1. While in reality, the motion is likely to be significantly more complex, this minimal model offers a straightforward way to incorporate the key stick-diffuse behaviour and make quantitative predictions. In particular, we can show that the MSD of a tracer particle exhibits a diffusive behaviour with the following effective diffusion constant *D*^eff^ = *p*_free_*k*_*B*_*T/*(6*πηa*) (see Methods). From this prediction, we expect the following relation to hold 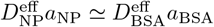, provided there is no significant size dependence in the probability to be in the free state *p*_free_. Using the radius of the nanoparticle tracers 100 nm and the Stokes radius of BSA 3.48 nm [62], we find the relation holds to a good approximation, supporting the scaling *D*^eff^ ∼ 1*/a*.

Chemical gradients produced by enzymes in the active droplets induce flows within the droplets upon interaction with surfaces, thereby activating the environment [63]. These flows may play a role in decreasing the rate at which particles are able to stick to current neighbors, mimicking a microscopic lubrication scenario. In the two-state model this corresponds to increasing *p*_free_ and therefore increasing the effective diffusion constant. From our results we can estimate that the activity-induced effects approximately halve the attachment rate of particles.

## MATERIALS

Polyethylene glycol (PEG) 4000 Da (A16151) was purchased from Alpha-Aesar. Potassium phosphate dibasic trihydrate (P5504), potassium phosphate monobasic (P5379), bovine serum albumine (BSA) (A7838), L-lactate dehydrogenase (LDH) (L1254),*β*- Nicotinamide adenine dinucleotide reduced disodium salt hydrate (NADH) (N8129), Pyruvate (P8524), Lactate assay Kit (MAK064), Yeast alcohol dehydrogenase (YADH) (A3263), *β*-Nicotinamide adenine dinucleotide hydrate, (NAD+) (N1511), 3-(Trimethoxysilyl)propylmethacrylate (440159), 2- Hydroxy-4’-(2-hydroxyyethoxy)-2 methylpropiophenone (410896), Poly(ethylene glycol) diacrylate (PEGDA) 700Da (455008) were purchased by Merck. Ethanol (14712-05), Acetaldehyde (00112-95) and Magnesium Chloride Hexahydrate (20909-55) were purchased from e-Nacalai Tesque. Alexa Fluor 488, 594 and 647 Microscale Protein Labeling Kit (A30006, A30008, A30009) were purchased from Thermo Fisher Scientific. Fluorescent polystyrene nanoparticles tracers of 0.2 *µ*m of diameter (FCDG003) were purchased from Bangs Laboratories.

## METHODS

### Droplet Preparation and Crowding Modulation

A stock solution of bovine serum albumin (300-350 mg/mL, around 30-35%) was prepared by mixing the appropriate amount of liophylized BSA in Milli-Q water or 0.1 M potassium phosphate buffer at pH 7.0 (the choice of the resuspension media is dependent on recipe used to prepare the droplets, see below). The concentration of the BSA stock solution was confirmed by measuring directly the absorbance (1:250 or 1:300 dilution) intensity at 280 nm using a UV-visible spectrophotometer (UV-1900 UV-Vis Spectrophotometer, Shimadzu) referring to *ε*_280_ = 43 824 M^*−*1^ cm^*−*1^ (https://web.expasy.org/protparam/).

PEG solution (600 mg*/*mL, 60% w/v) was prepared by mixing and heating at (40 ^*°*^C for 20 minutes) the appropriate amount of PEG 4000 and Milli-Q water or buffer KP 0.1 M pH 7.0 (see below). At this point, droplet resuspensions were prepared using different recipes depending on the desired amount of BSA inside the droplet.

The droplet resuspension was prepared using a final target concentration of 30 mg*/*mL BSA (3% w/v), 230 mg*/*mL of PEG 4000 (23% w/v), 200 mM KCl and a final concentration of 100 mM KP buffer at pH 7.0 in a microcentrifuge tube. PEG has been added as last component during droplet preparation. Additional components, such as enzymes, cofactors, ions and other molecules were added to the tube if needed. The tube was gentle shaken to promote the mixing. When required the droplet resuspension was centrifuged for 60 minutes at 16 900 g and the two phases separated by pipetting.

In this standard condition, the BSA within the droplets phase is equal to 45 % w/v, which corresponds to a concentration range between 6.5-7 mM inside of the droplet. In order to decrease the concentration of BSA inside the droplet we have used different recipes (by modulating initial BSA and PEG concentration) to prepare the droplet resuspension. Please not that in this case we have resuspended the stock solution of BSA and PEG in buffer KP 0.1 M pH 7.0 rather than MilliQ water to avoid changes in the final ionic strength of the droplets resuspension and keep the final buffer concentration 0.1 M pH 7.0. In detail, by using a final target concentration of 90 mg*/*mL BSA (9% w/v), 170 mg*/*mL PEG 4000 (17% w/v), 200 mM KCl and a final concentration of 100 mM KP buffer at pH 7.0 the droplet phase has a BSA content of 37% w/v, corresponding to a concentration range of 5.5- 5.8 mM BSA. The concentration of the BSA stock solution was confirmed by measuring directly the absorbance (1:250 or 1:300 dilution) intensity at 280 nm using a UV-visible spectrophotometer (UV-1900 UV-Vis Spectrophotometer, Shimadzu) referring to *ε*_280_ = 43 824 M^*−*1^ cm^*−*1^ (https://web.expasy.org/protparam/).

Lastly, using a final target concentration of 140 mg*/*mL BSA (14% w/v), 150 mg*/*mL PEG 4000 (15% w/v), 200 mM KCl and a final concentration of 100 mM KP buffer at pH 7.0, the droplet phase has a BSA content of 32% w/v, corresponding to a concentration range of 4.5-5.0 mM BSA. More than 20 measurements have been performed for each droplet phase in all the reported conditions. Please note that the uncertainties of the BSA and PEG 4000 in the above recipes are about 0.5-1% (w/v) and the uncertainties of the BSA concentration in the droplet phases is 1.5-3 (w/v). The volume fraction of the droplet phase has been measured as reported here [11]. In detail, droplets containing 45% w/v BSA show a droplets volume fraction between 2-5 %, droplets containing 37% w/v BSA show a droplets volume fraction between 9-14% and droplets containing 32% w/v BSA show a droplets volume fraction between 18-24%.

### Enzyme Protein Labeling

L-lactate dehydrogenase (LDH) and Alcohol dehydrogenase (ADH) were tagged using Alexa Fluor 594 and 647 Microscale Protein Labeling Kit respectively, using the manufacturer’s protocol but repeating the dialysis step three times to remove the free dye in the protein solution. Tagged proteins were further purified using membrane dialysis to reduce the free probe in solution. The presence of the free dye and the integrity of the proteins after labeling has been checked using SDS-PAGE (data not shown). To determine the protein concentration of LDH and ADH we used *ε*_280_ = 43 824 M^*−*1^ cm^*−*1^ and *ε*_280_ = 47 370 M^*−*1^ cm^*−*1^ respectively (https://web.expasy.org/protparam/).

### Confocal microscopy

The fluorescence images were acquired and analyzed using a Spinning Disk Confocal (Nikon, Andor CSU) microscope with × 63 oil immersion lens. Confocal images were acquired for more than 5 independent experiments with similar results. The images collected were analyzed using ImageJ.

### Activity measurements of LDH inside droplet resuspension

#### Kinetic parameters of L-lactate dehydrogenase

The kinetic parameters of LDH (*k*_cat_ and *K*_M_) have been calculated in all the experimental conditions (45%, 37% and 32% w/v of BSA in the droplet phase) using 2 mM NADH and pyruvate concentration ranging from 0.1 to 1 mM. As internal controls, we have used several dilutions (500-fold to 1000-fold) of LDH-containing droplets and several LDH concentrations within the droplets to calculate the kinetic parameters and we have observed no substantial changes in the kinetic values. The final concentration of LDH in the droplet resuspension ranged from 0.5 µM to 3.3 µM, due to the different final droplets volume fraction obtained by preparing the droplets with different recipes (32%, 37% and 45% w/v), which led to a different concentration of LDH inside the droplets even when adding the same amount of free enzyme in the mix before droplet preparation. However, as mentioned above, in the LDH concentration range between 0.5 µM to 3.3 µM we have observed no significant changes of the kinetic parameters (). At each time point for each substrate concentration, the reaction mix was stopped by decreasing the pH, by adding 5 µL of trichloroacetic acid (TCA) at a final concentration of 10% (v/v, 45 µL of reaction mix + 5 µL of TCA 100%). After appropriate mixing, we stored the stopped reaction mix in ice protected from the light for at least 10 minutes. The 50 µL of stopped reaction mix has been then centrifuged for 20 minutes at 16 900 g at room temperature to emove the unfolded proteins and clarified the stopped reaction mix before lactate determination.

The lactate concentration was determined by using the Lactate assay Kit using a Multiskan SkyHigh Microplate Reader (Thermo Fisher Scientific) as previously reported [11]. To calculate the *k*_cat_ value the absorbance values were converted into nmol of product and then to reaction rate values using the nmol of the enzyme used at the time of the assay (as already reported here [11]). The reaction rate values at each pyruvate concentration were fitted using the Michaelis-Menten equation [64]. All the experiments have been performed in triplicate (n = 3) and are reported as mean ± standard deviation. Data analysis was performed using GraphPad Prism 10.4.1.

#### Partitioning of LDH

Activity measurements have been performed as explained here with modifications [11, 65]. Briefly, the partitioning of LDH (ranged from 0.5 µM to 3.3 µM) inside the droplet containing BSA at 45%, 37% and 32% (w/v). We measured the activity of the enzyme in the supernatant phase that remained after droplets were removed through a centrifugation step. The assay was performed in presence of 10 mM NADH and using a fixed pyruvate concentration equal to 0.8 mM. We stopped and treated the reaction mix (using a final concentration of 10% TCA) and quantified the lactate produced as described above.

### Activity measurements of ADH inside droplet resuspension

The kinetic parameters of ADH (*k*_cat_ and *K*_M_) in the droplet resuspension have been calculated in all the experimental conditions in presence of 10 mM NAD+ and using ethanol as substrate, modulating its concentration from 15 to 1000 mM. Droplets solution containing partitioned 0.5 µM to 3.3 µM ADH were assayed for different time points and several substrate concentrations monitoring the formation of NADH at 340 nm. As already reported for LDH above, the final concentration of ADH in the droplet resuspension ranged from 0.5 µM to 3.3 µM, due to the different volume fraction of the droplets prepared with the different recipes (32%, 37% and 45% w/v), which led to a different concentration of ADH inside the droplets even adding the same amount of free enzyme in the mix before droplet preparation. Briefly, 0.4 mL of droplets containing partitioned ADH were prepared and 1 µL was withdrawn and diluted 500-fold in the supernatant containing different substrate concentrations. At each time point, 200 µL of the reaction mix was stopped by heating the sample at 100 ^*°*^C inside a water bath for 2 minutes and then storing in ice for 5 minutes. Note that in our system we did not observe any NADH degradation by heating the reaction mixture for 2 minutes (data not shown). Then, the samples were centrifuged at room temperature for 30 minutes at 16 900 g to clarify the reaction mix by removing the unfolded protein for NADH quantification. The NADH quantification was performed by placing 100 µL of the stopped reaction mix in a 96-well plate and the absorbance at 340 nm was measured. An NADH calibration curve ranging from 0 to 500 µM was made in all the experimental conditions, using high purity NADH. To calculate the *k*_cat_ value the absorbance values were converted into nmol of product and then to reaction rate values using the nmol of the enzyme used the time of the assay (nmol of product/nmol of enzyme/time expresssed in seconds). The reaction rate values at each ethanol concentration were fitted using the Michaelis-Menten equation [64]. All the experiments have been performed at least in triplicate (n = 3) and are reported as mean ± standard deviation. Data analysis was performed using GraphPad Prism 10.4.1.

The activity of ADH can also be measured using the continuous assay by monitoring the NADH formation over time at 340 nm. However, the linearity of NADH formation is often confused with the cloudy aspect of the solution typical of phase separated systems which makes difficult to clearly follow the NADH signal. The partitioning of ADH has been evaluated inside the droplet containing 45%, 37% and 32% (w/v) BSA and using 0.5-2 µM ADH partitioned in the droplets. We measured the activity of the enzyme in the supernatant phase that remained after droplets were removed through a centrifugation step and we compared the activity with that obtained in the droplet resuspension. The assay was performed in presence of 10 mM NAD+ and using a fixed ethanol concentration equal to 100 mM. We stopped, treated the reaction mix and measured the enzymatic activity as described above.

### FRAP experiments

The fluorescence recovery after photobleaching (FRAP) experiments were carried out using a laser confocal system Nikon A1R System on a Nikon TiE2 inverted microscope using an 100x oil objective (100 × /1.49 Oil Apo TIRF). The 488 nm laser line was used for both fluorescence photobleaching and detection. The fluorescence decay and recovery were measured using a GaAsP detector (Nikon DUB4). All the experiments were designed and carried out using Nikon NIS Elements software. The data were exported as single Excel file to be then processed in a MATLAB-based routine. The recovery time (seconds) was extracted from the fitting of normalized time-dependent intensity curves using a least squares regression. Imaging array and pixel resolution were optimized to obtain a sampling ratio between 4 and 15 frames per second depending on how fast the kinetics were considered in this study. An average bleaching spot of 1.5 microns in diameter was obtained throughout the experiments with a bleaching duration ranging between 115 and 215 msec. The temperature was kept constant at 298 K. The imaging settings were adjusted such that at least 500-1000 images could be acquired without an overall bleaching of more than 10% of the initial signal.

The bleaching experiments consisted of three phases starting with the acquisition of 20 - 100 pre-bleach images to determine the steady state pre-bleach value of the fluorescence signal and to ensure that the focal plane was maintained during the imaging. A reasonably short bleach spot was then applied approximately in the middle of the target droplet. The depth and precision of the bleached region were determined by a simple line measurement across the bleached region. The bleach duration was minimized but a bleaching efficiency of no less than 85% was maintained. Typically, 10-15 droplets from the same batch were imaged for each experiment (total duration *<*45 min to prevent artifacts due to the aging of the droplets) with and without substrate. The droplets were freshly prepared before each set of measurements. The volume containing droplets were seeded in a small chamber (200 microliters) glued on a coated fluorescence microscopy suitable coverslip. Different concentrations of substrate were added to the solution depending on the experimental condition set-up. After a short settling time the kinetic measurements were initiated (generally 2-4 minutes).

To extract diffusion constants from the FRAP data, we numerically solved the 2D diffusion equation of a normalised concentration of fluorescent particles. The initial condition was a 1.5 µm diameter circular patch of low concentration surrounded by a bulk of high concentration, matching the experimental conditions. As can be seen in Fig. 3**a**, this model system accurately reflects the dynamics seen in experiment. The inset of Fig. 3**b** shows theoretically calculated intensity recovery curves on top of experimentally measured ones, again highlighting that the experimental system is well represented by the the-oretical model. Using the model we calculated FRAP half-lives for different diffusion constants to obtain the curve shown in Fig. 3**b**. We then inverted this curve to obtain diffusion constants given experimentally measured FRAP half-lives. In Fig. 3**b** we also show a half-life curve for systems with a flow velocity of the order that we measure in tracking experiments. There is no significant difference in the curve to having no flow, since at this scale diffusion dominates. However as discussed, the presence of flow indirectly appears in the FRAP results since it affects the diffusion constant. For more details on the FRAP analysis, please see the Supplementary Information.

### Nanoparticle tracking experiments

Green fluorescent polystyrene nanoparticles tracers of 0.2 µm average diameter were added to the droplet suspension, at final concentration of 0.0005-0.0008% solid content. The green fluorescent nanoparticles displayed good partitioning in the droplet phase, with only marginal aggregation visible and by using the highest nanoparticles concentration. We recorder timelapses of fluorescent tracers with a spinning disc confocal microscope (Nikon) controlled by 3i software (time interval = 2 s) and tracked their motion in 2D. The Python package trackpy [66] was used to identify and track nanoparticles in the videos and to subsequently analyse their motion. In total six active and six inactive droplets were analysed. We prioritised the quality of trajectories over the quantity of trajectories per droplet, applying the necessary thresholds for feature size and brightness and filtering out any trajectories shorter than 200s. This resulted in around 10-25 independent trajectories per 0 mM pyruvate droplet, and around 20-50 trajectories per 1 mM pyruvate droplet. The trajectory duration distribution was spread from 200 s up to around 1000 s relatively uniformly. This procedure ensured good quality MSDs for the timescales we used (up to 140 s). For more details on the tracking procedure, please see the Supplementary Information.

### Theoretical calculation of the flow structure

A sustained chemical gradient in the vicinity of a surface or an interface can lead to the emergence of a sustained flow, which has a circulating structure due to the incompressibility of the solvent, and can thus activate the interior of a chemically active droplet [63]. To solve for the flow profile, we need to connect the concentration profile of the chemical *c* and the density profile of the enzymes *ρ* to the flow velocity of the solvent ***v*** via an induced surface-slip phoretic velocity boundary condition near the surface [67]. Solving the relevant equations (see SI), we obtain the emergent velocity flow inside and outside the droplet. The exact solution, which is azimuthally symmetric, is expressed in spherical coordinates with *r* being the distance from the centre of the hemisphere and *θ* being the polar angle. The region inside (outside) the droplet is *r* ≤*R* (r ≥ R) where *R* is the radius of the droplet.

The solution for the enzyme density profile inside the droplet is given by

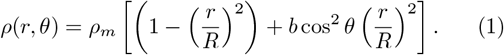

By definition, the density vanishes for r ≥ R. The value of the dimensionless coefficient *b* – which is formally needed for the purposes of consistency in terms of the boundary conditions – is found to be sufficiently small (*b* ≃ 10^*−*8^) to justify neglecting it for practical purposes. This density profile is shown in Figs. 3**g**-**h** as a heatmap.

The radial and polar component of velocity flow inside the droplet are found as

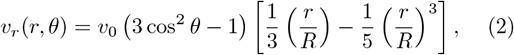

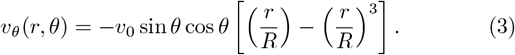

Here

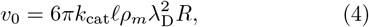

sets the relevant velocity scale in the problem, where *ρ*_*m*_ is the maximum enzyme density scale, *λ*_D_ is the Derjaguin length [63], and *𝓁* is the radius of the solute particles. Using typical values for the parameters (λ_D_ ∼ 1 nm, *𝓁* ∼ 1 nm, *R* ∼ 10 µm, *k*_cat_ ∼ 10^3^ s^*−*1^, and *ρ*_m_ ≃ 1 µM), we find the value of *v*_0_ ∼ 0.1 µm s^*−*1^. This flow profile is shown in Figs. 3**g**-**h** as streamline plots and in Figs. 3**h**-**I** as velocity profiles, as well as in Fig. 1 as a 3D streamline plot. This calculation allows us to predict the strength of the velocity in different horizontal planes inside the bubble as shown in the Fig. 3**h**. We note that we have chosen to demonstrate the existence of this mechanism at the presence of a single concentration field for simplicity, but we can easily the calculation to include multiple concentrations due to the linearity of the equations [63].

### Diffusion in a heterogeneous environment

We can characterize the stochastic trajectory of a tracer particle, described by position ***r***(*t*), in an environment where it frequently experiences sticky (i.e. highly attractive and localized) interactions and consequently slows down, using a Langevin dynamics with a random time-dependent diffusion coefficient *D*(*t*), namely

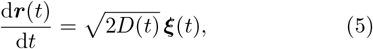

where ***ξ***(*t*) is a Gaussian white noise term of zero mean and unit strength, namely ⟨ *ξ*_*i*_(*t*) ⟩ = 0 and ⟨ *ξ*_*i*_(*t*)*ξ*_*j*_(*t*^*′*^) ⟩ = *δ*_*ij*_*δ*(*t* −*t*^*′*^) with *i, j* {1, · · ·, *d*} in *d* dimensions. The apparent, or observable, mean-squared displacement can then be calculated as

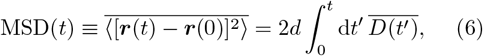

where the over-line denotes quench-averaging over the distribution of diffusion coefficients. We now use a simple two-state description, in which the tracer is either *free* with probability *p*_free_, where the diffusion coefficient is *D*_free_ = *D*_0_, given by the Stokes-Einstein relation corresponding to the free liquid environment *D*_0_ = *k*_B_*T/*(6*πηa*) (with *η* being the viscosity of the solvent, *a* being the radius of the particle, and *k*_B_*T* being the thermal energy), or *stuck* with probability *p*_stuck_ = 1 − *p*_free_, where the diffusion coefficient is *D*_stuck_ = 0. This yields 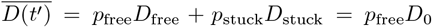, and a resulting effective or apparent diffusion coefficient given as

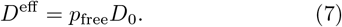

### MD simulations

The X-ray structure of rabbit muscle LDH (see Fig. S11) in complex with NADH and oxamate (a pyruvate analog) (PDB code: 3H3F) [68] was employed for our molecular dynamics (MD) simulations. Pyruvates were prepared at the active sites by converting the amino group of oxamate to a methyl group. All histidine residues were treated as neutral except for protonated His192. One LDH ternary complex in its tetrameric form was solvated in a cubic water box with dimensions of 136 Å × 136 Å × 136 Å, and 150 mM KCl was added to construct a dilute condition. For consistency with experimental conditions, bovine serum albumin (BSA, PDB code: 4F5S) [69] was used as a crowder to build a crowding condition. Two, four, or eight BSAs were included in a simulation box (termed as 2bsa, 4bsa, or 8bsa, respectively) to mimic crowding conditions. Furthermore, ligand-free (apo) LDH was prepared both in dilute and crowding conditions. These systems were prepared in the same way, except that four NADH and pyruvate molecules were randomly distributed in the solution. Following the previous work [70], Amber ff99SB-ILDN [71, 72] and TIP3P [73] force fields were used for protein and water, respectively. The parameters for NADH and pyruvate were derived by GAFF2 force field with AM1-BCC [74]. The solvation box was generated for each system by tleap of AmberTools [75]. During MD simulations, the covalent bonds involving hydrogens in proteins and water molecules were constrained through SHAKE and SETTLE algorithms, respectively [76, 77]. The long-range electrostatic interaction was calculated using the particle mesh Ewald (PME) method [78–80]. BUSSI thermostat was applied to maintain the NVT ensemble [81]. The system was propagated using the r-RESPA multiple time step method with a time step of 3.5 fs for fast motions and 7.0 fs for slow motions [82]. The optimized hydrogen mass repartitioning (HMR) technique and a group-based temperature/pressure approach (group T/P) were employed to make the simulation stable [83, 84]. For each system, we carried out two independent simulations. In order to eliminate the influence of the initial configuration, we prepared two distinct initial structures for every crowded system by randomly rearranging the placement of crowders (BSAs) around LDH. To alleviate steric clashes, we performed four consecutive rounds of energy minimization, consisting of 20000 steps each, gradually reducing the positional restraints of protein and crowder molecules. Following a 100 ps heating process from 0 K to 300 K, a series of short NVT equilibrations at 300 K were conducted to allow the system to relax. Throughout the equilibration, the positional restraints on the backbone of Ca atoms of LDH and the heavy atoms of substrates were gradually removed. Subsequently, we performed 520 ns conventional MD (cMD) simulations in the NVT ensemble at 300 K as the production run. Flat-bottom distance restraints were introduced between an oxygen atom of pyruvate (O1) and a nitrogen atom of His192 (NE2) with a weak force constant, 1.0 kcal · mol^*−*1^Å^*−*2^. The restraints were set to be effective when the O1-NE2 distance was greater than 4 Å, thereby ensuring that pyruvates remained stably bound to their respective sites in every subunit.

### QM/MM calculation

The holo form of LDH tetramer, in the presence of NADH and pyruvate, was solvated in a cubic water box with dimensions of 114 Å in each direction with 150 mM KCL. Following multiple steps of energy minimization, the system was equilibrated through MD simulations in the NVT ensemble at 300 K with the PBC condition. During the equilibration process, the positional restraints of the LDH backbone and sidechain were gradually reduced while retaining those on substrates. Then, the cubic water box was converted into four water spheres with a radius of 40 Å centered at the C_*α*_ atom of Ser160 of each subunit (Fig. S12). This cluster system (without periodic boundary conditions) was used to perform QM/MM calculations. The QM region was set to 39 atoms located at the active site, i.e., the pyruvate molecule, the nicotinamide ring of NADH, and the side chain of His192 (Fig. S12**b**), whereas the surrounding proteins and solutions were treated by the AMBER force field. Subsequently, the pyruvate of chain A or chain B (open or closed form, respectively) was relaxed through a series of QM/MM-MD calculations by reducing the positional restraints on pyruvate while keeping weak restraints on the Ca atoms of LDH. The DFTB3 [85] method was used for the QM calculations. The equilibrations were performed in the NVT ensemble controlled at 300 K using the BUSSI thermostat64. Velocity Verlet (VVER) [86] integrator was used with the time step 2.0 fs. The switching and cut-off distance of the electrostatic interaction was set to 16.0 and 18.0 Å, respectively.

The string method was employed to predict the minimum energy pathways (MEPs) of the enzyme reaction of LDH in both open and closed forms [87, 88]. Prior to the string simulation, the conformations of the reactant (His192H+, pyruvate, and NADH) and product (His192, lactate, and NAD+) were calculated for each form. The structure of the reactant was obtained by QM/MM energy-minimization of the active site of chain A or B (open or closed form, respectively) at the level of density function theory (DFT) with B3LYP-D3 [89–91] functionals and aug-cc-pVDZ basis sets [92]. Then, the geometry optimization was carried out in two steps to generate the product. In the first step, the optimization was carried out with distance restraints added to dissociating bonds (the reference distances set to 1.7–2.0 Å) and newly formed bonds (the reference distances set to 1.0 Å). In the second step, structural refinement was carried out without any restraints. The initial pathway of string simulations was prepared by linear interpolation of the reactant and product in terms of Cartesian coordinates. The pathway was divided into 16 evenly spaced images. The active atoms in a path search were the same as the QM region. The step size was set to *δt*=0.5 fs. The energy minimization was performed using a limited memory version of Broyden-Fletcher-Goldfarb-Shanno (L-BFGS-B) [93, 94] and a macro/micro-iteration scheme [95]. The IAO charges [96] were derived from the electron density obtained in B3LYP-D3 calculations. Convergence thresholds of the energy profile and the path length were set to 0.01 kcal · mol^*−*1^ and 0.01 Å, respectively. The string simulations of both closed and open forms showed a smooth convergence (Fig. S13).

To derive the free energy profile of the enzyme reaction, we carried out umbrella sampling (US) calculations along the MEP predicted by the string method. The bond distances that undergo significant changes during the reaction (*r*_1_ – *r*_7_ depicted in Fig. S14) were selected as the collective variables (CVs). The CVs were restrained with a harmonic force constant of 200 kcal · mol^*−*1^Å^*−*2^. 28 and 32 windows were used for the US simulations of open and closed forms, respectively, to enable sufficient overlap of the probability distribution between neighboring windows. The windows were generated in an equidistant manner along the MEPs in the CV space. For each window, an equilibration MD was carried out for 1 ps using DFTB3/MM, followed by a production run for 2.5 ps using B3LYP-D3/MM. The integration time step was set to *δt* = 0.5 fs, and the hydrogen atoms involved in the reaction were unconstrained. The simulations were performed at T =300 K in the NVT ensemble. To avoid the large conformational fluctuation of LDH, weak positional restraints were applied to the C_*α*_ atoms of LDH (the force constant was set to 0.5 kcal · mol^*−*1^Å^*−*2^). The multistate Bennett acceptance ratio (MBAR) method [97] was used to reweight the trajectory and 1D- and 2D-PMF were obtained from weighted histograms.

### Software

All the MD simulations and QM/MM calculations were carried out using GENESIS [98, 99] software. The QM calculations were conducted using QSimulate-QM [100]. The use of a highly parallelized QM/MM program combining GENESIS and QSimulate-QM, which can achieve tens of picoseconds of DFT/MM-MD calculation per day [101], was critical for the present study. The molecular visualization program PyMOL [102] was employed to generate the structure graphs. Matplotlib [103] was utilized for plot preparation. MDtraj [104] and GENESIS analysis tools were utilized to analyze the trajectories. The trackpy Python package [66] was used to track and analyze the nanoparticle motion.

## CODE AVAILABILITY

The algorithms for the codes supporting the main findings of this study are available in the paper and its Supplementary Information. Any additional information concerning the code can be made available upon request.

## ACKNOWLEDGEMENTS

The research was supported by the Okinawa Institute of Science and Technology Graduate University (OIST) with subsidy funding to P.L. from the Cabinet Office, Government of Japan. This work was supported in part by MEXT JSPS Kakenhi (grant number 19H05645, 21H05249 (to Y.S.), 22H04761, 24K01446 (to K.Y.),24K01995 (to P.L.) 22K15065 (to M.D.)), RIKEN pioneering projects “Biology of Intracellular Environments”, and “Glycolipidologue Initiative” (to Y.S.), MEXT program for promoting research on the supercomputer Fugaku (JPMXP1020200101), and MEXT program for Big-data-driven bio/synthetic polymer science to create absolutely circular materials (JPMXP1122714694) and Data-Driven Research Methods Development and Materials Innovation Led by Computational Materials Science (JPMXP1020230327) (to Y.S.). The computer resources are provided by the HPCI system research project (Project ID: hp230043, and hp240057). We are grateful for the help provided by imaging section at OIST; in particular we thank Paolo Barzaghi for the FRAP experiments and analysis. R.G. acknowledges support from the Max Planck School Matter to Life and the MaxSynBio Consortium which are jointly funded by the Federal Ministry of Education and Research (BMBF) of Germany and the Max Planck Society.

## AUTHOR CONTRIBUTIONS

M.D. and P.L. conceived the project and designed the experiments. M.D. performed the in-vitro experiments. M.D. and P.L. analyzed the experimental data. W.R., K.Y., and Y.S. designed the MD simulations and QM/MM calculations. W.R. and K.Y. performed the calculations. W.R., K.Y., and Y.S. analyzed the trajectory data. J.M. and R.G. performed the FRAP diffusion coefficient analysis and the nanoparticle tracking analysis. M.C., J.M., and R.G. performed the calculation of the velocity flow field. All authors wrote the manuscript. P.L., Y.S., and R.G. supervised the project.

