## Supplementary Information for "Enzymes can activate and mobilize the cytoplasmic environment across scales"

(Dated: January 28, 2025)

#### CONTENTS

I. Supplementary Figures 2

II. FRAP analysis 19
A. Theoretical FRAP model 19
B. Numerical integration 19

III. Nanoparticle tracking 20
A. Obtaining trajectories 20
B. Calculating MSDs and extracting parameters 20

IV. Derivation of the induced velocity flow field 20
A. Hydrodynamics 20
B. Connecting flow velocity and concentration 21
C. Projected velocity 23

References 23

*
†
‡

### I. SUPPLEMENTARY FIGURES

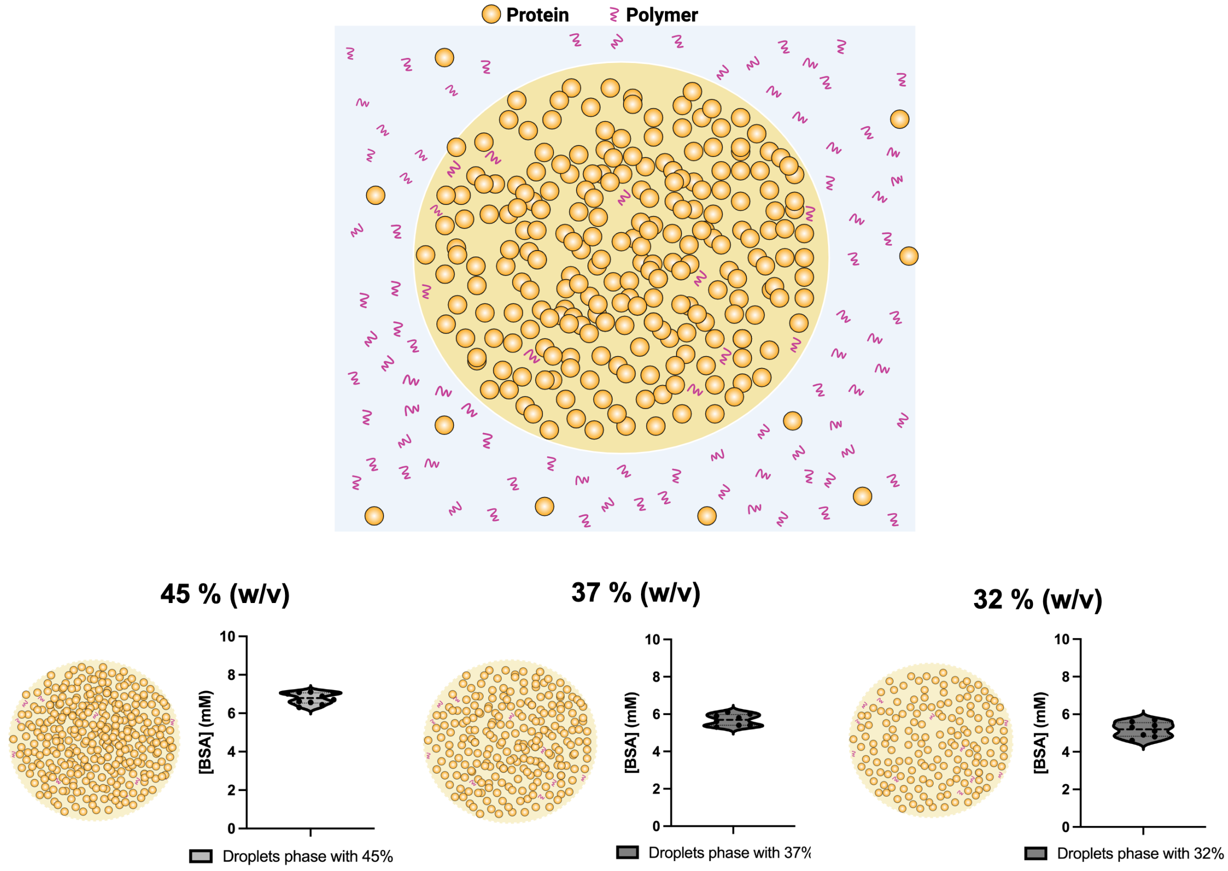


FIG. S1. **Schema of the droplets system and quantification of the protein concentration (BSA) within the droplets phase. The droplets have been prepared by using different BSA concentration (45%, 37% and 32% in the droplet phase expressed as w/v).** Schema of the droplets system and spectrophotometric quantification of the protein concentration within the droplets (45%, 37% and 32% BSA w/v) prepared by mixing different concentrations of BSA and PEG. The BSA concentration has been calculated by measuring the absorbance at 280 nm as reported in the Material and Methods section.

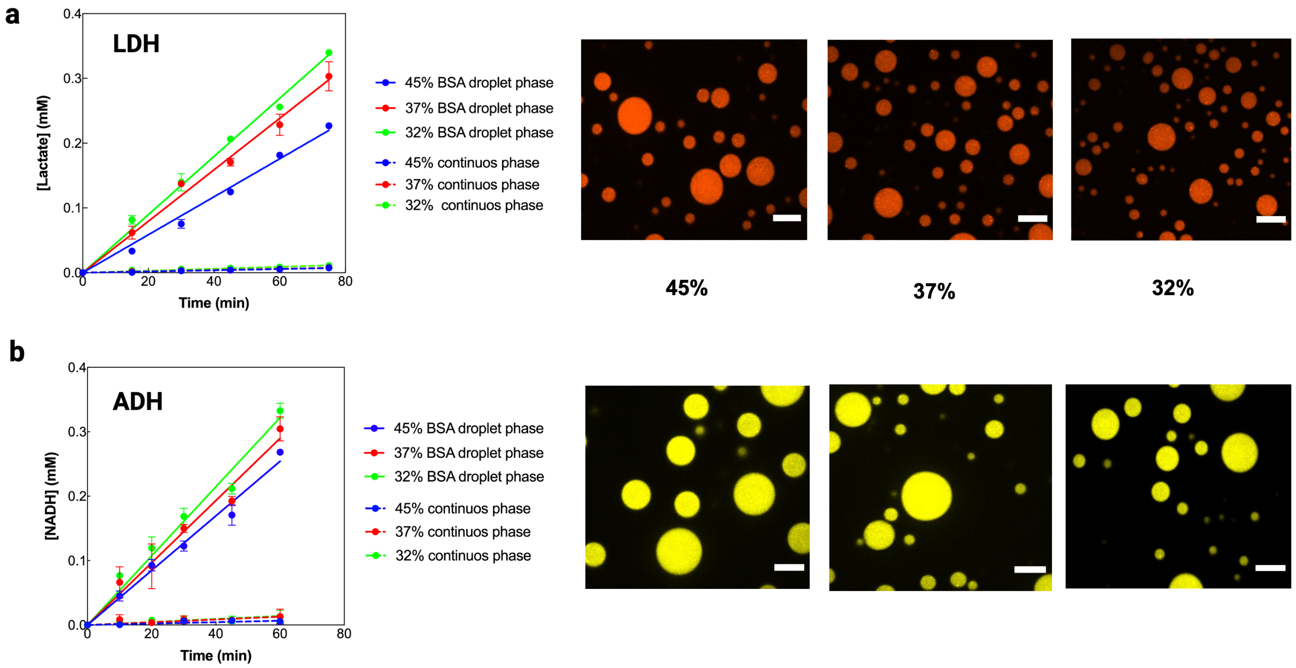


FIG. S2. **Evaluation of the enzymes partitioning within the droplets containing different protein concentration.** (a), Confocal microscopy images of Alexa Fluor 594-labeled LDH (red droplets, panel a) and Alexa Fluor 488-labeled ADH (panel b) partitioned inside liquid-liquid phase separated droplets containing different BSA concentration within the dense (droplets phase) (top to bottom (BSA w/v): 45%, 37% and 32%). Confocal images were acquired for more than 5 measurements LDH activity within 45%, 37% and 32% w/v of BSA in the droplets phase is highlighted by the colored dots and lines which represent the amount of lactate produced by the enzymes over time. The dots and dashed lines represent the amount of product lactate measured in the continuous phases obtained after centrifugation and removal of the droplets phase. (b), ADH activity within the droplets (45%, 37% and 32% w/v BSA) is highlighted by the colored dots lines which represent the amount of acetaldehyde produced over time. The dots and dashed lines represent the amount of NADH produced in the continuous phase, which has obtained after centrifugation and removal of the droplets phase. Both the enzymes within the droplets were used at the final concentration of 3.3 μM.

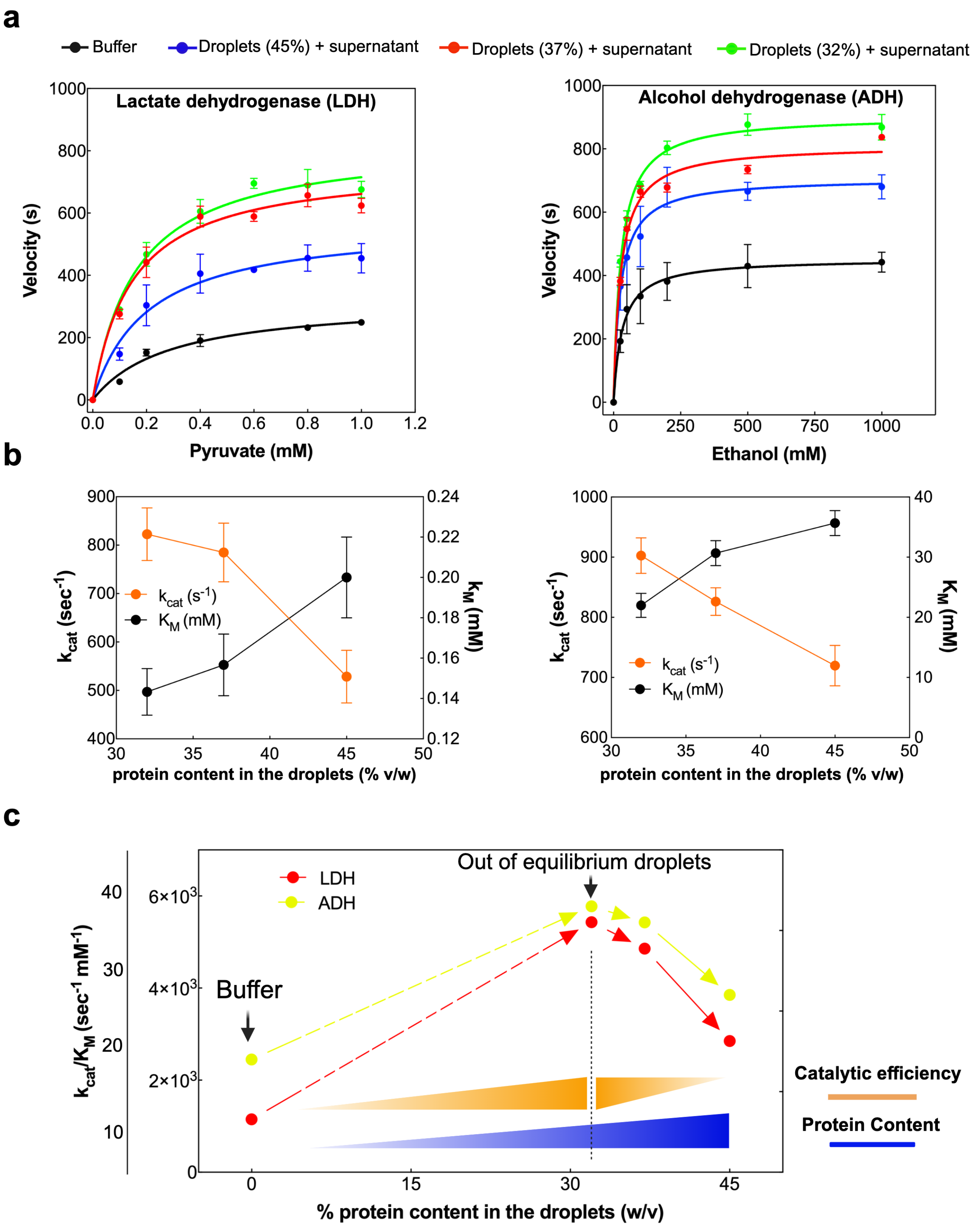


FIG. S3. **Michaelis-Menten graphs and kinetic parameters expressed as $k_{\text{cat}}$, $K_{\text{M}}$ and $k_{cat}/K_m$ in the droplets containing different protein concentration.** (a), Michaelis-Menten plot of LDH and ADH obtained in dilute conditions (no droplets, black line and dots) and with the enzymes partitioned within droplets containing different protein concentration (45% BSA (w/v), blue line and dots, 37% BSA (w/v), red line and dots and 32% BSA (w/v), green line and dots). (b), Trend of the $k_{\text{cat}}$ and $K_{\text{M}}$ values of LDH and ADH plotted versus droplets containing different protein concentration (% BSA (w/v)). (c), Schema representing the trend of the $k_{cat}/K_m$ values for LDH and ADH, highlighting the difference between the measured values of the $k_{cat}/K_m$ in diluted (buffer) and crowded conditions (droplets) and the variation of the $k_{cat}/K_m$ values in droplets containing different protein concentration.

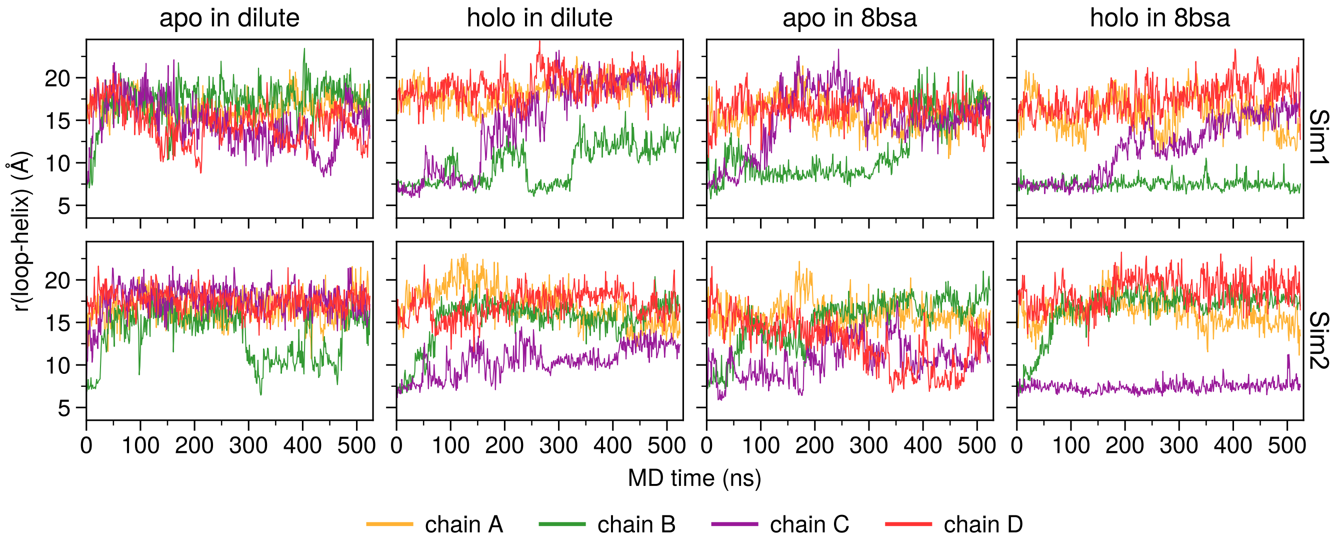


FIG. S4. **The time evolution of the distance between the active site loop and its contacting helix for apo and holo states in dilute and crowded (8bsa) conditions.** The LDH monomer in each chain is depicted in different colors. The initial configuration of the LDH tetramer had two monomers (chain A and chain D) in an open form and the other two monomers (chain B and chain C) in a closed form. The data from two MD simulations (Sim1 and Sim2) are presented in the first and second rows, respectively.

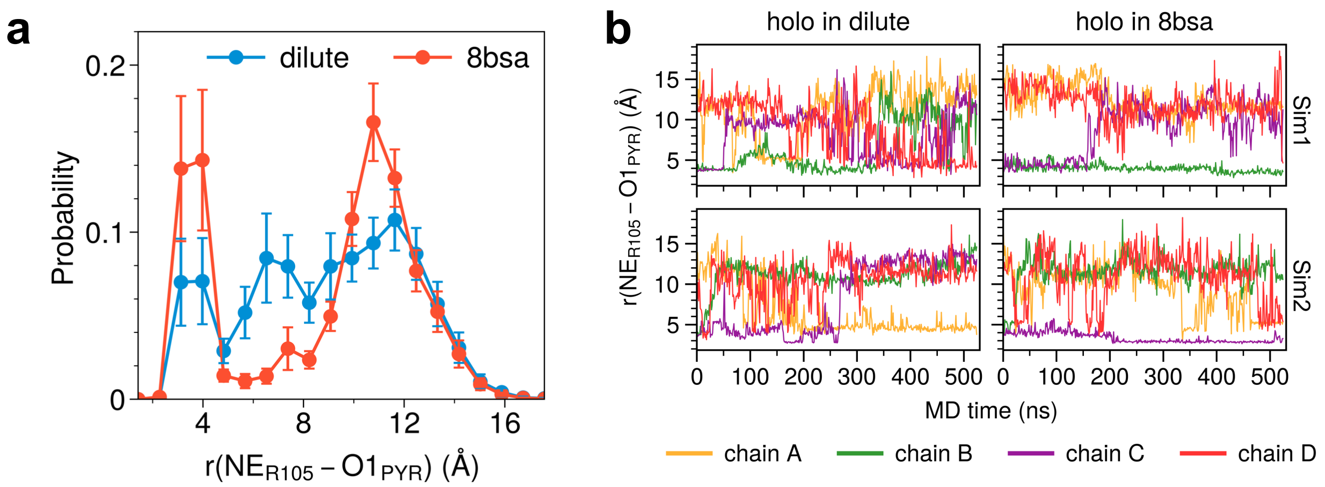


FIG. S5. **The distance between Arg105 and and pyruvate.** (A) A histogram of the distance between a nitrogen atom of Arg105 (NER105) and an oxygen atom of pyruvate (O1PYR) in dilute and crowded conditions. (B) The time evolution of the NER105-O1PYR distance in each LDH monomer in dilute and crowded conditions from two independent simulations.

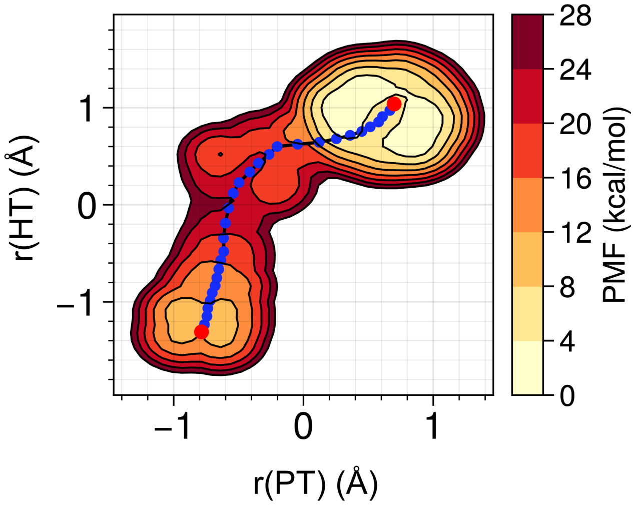


FIG. S6. **Two-dimensional PMF in terms of proton transfer (PT) and hydride transfter (HT) coordinates in the open state.**

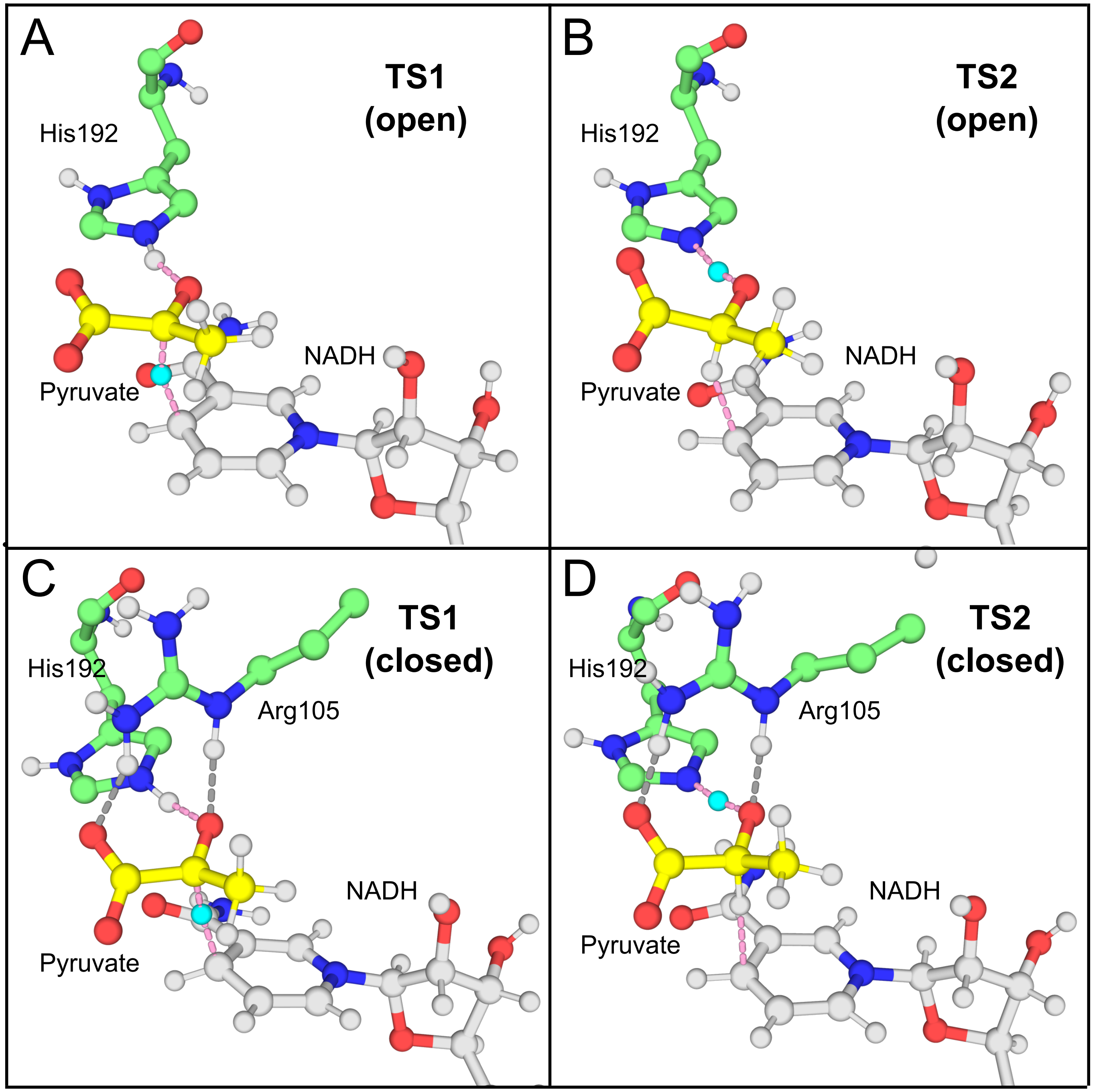


FIG. S7. **Typical configurations of transition states of HT (TS1) and PT (TS2) reactions in the closed and open forms of LDH monomers, respectively.** The transferring hydride and proton are depicted as cyan spheres, while the breaking and associating bonds are represented by pink dashed lines. Interactions between Arg105 and pyruvate are indicated by grey dashed lines.

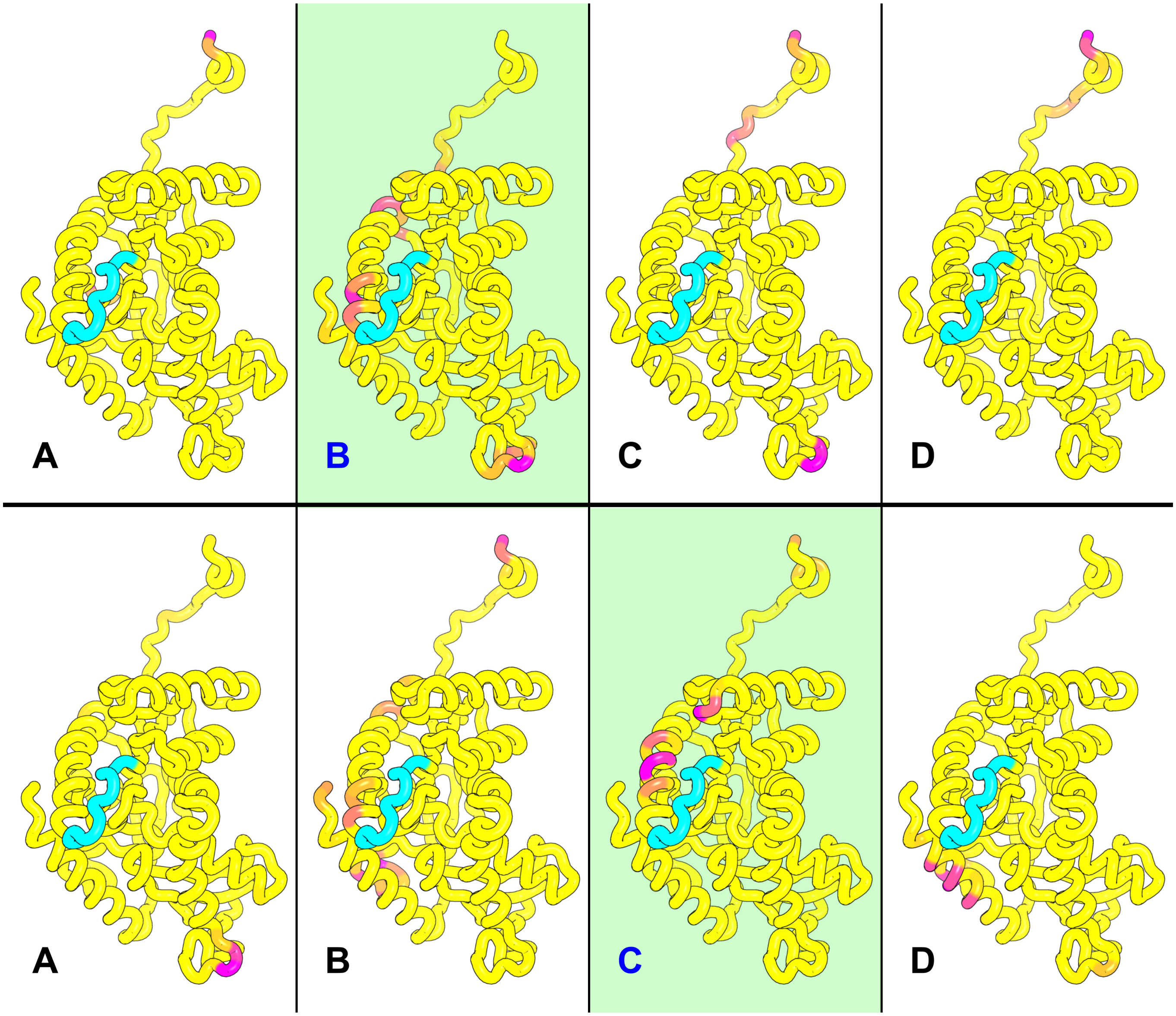


FIG. S8. **The probability of amino acids in each LDH monomer (chain A, B, C, and D) forming contact with BSA.** The higher probabilities are indicated by deeper shades of pink. The active site loop is depicted in cyan. The stable closed monomer is shaded in green. Data from two independent simulations are presented in the top and bottom panels separately.

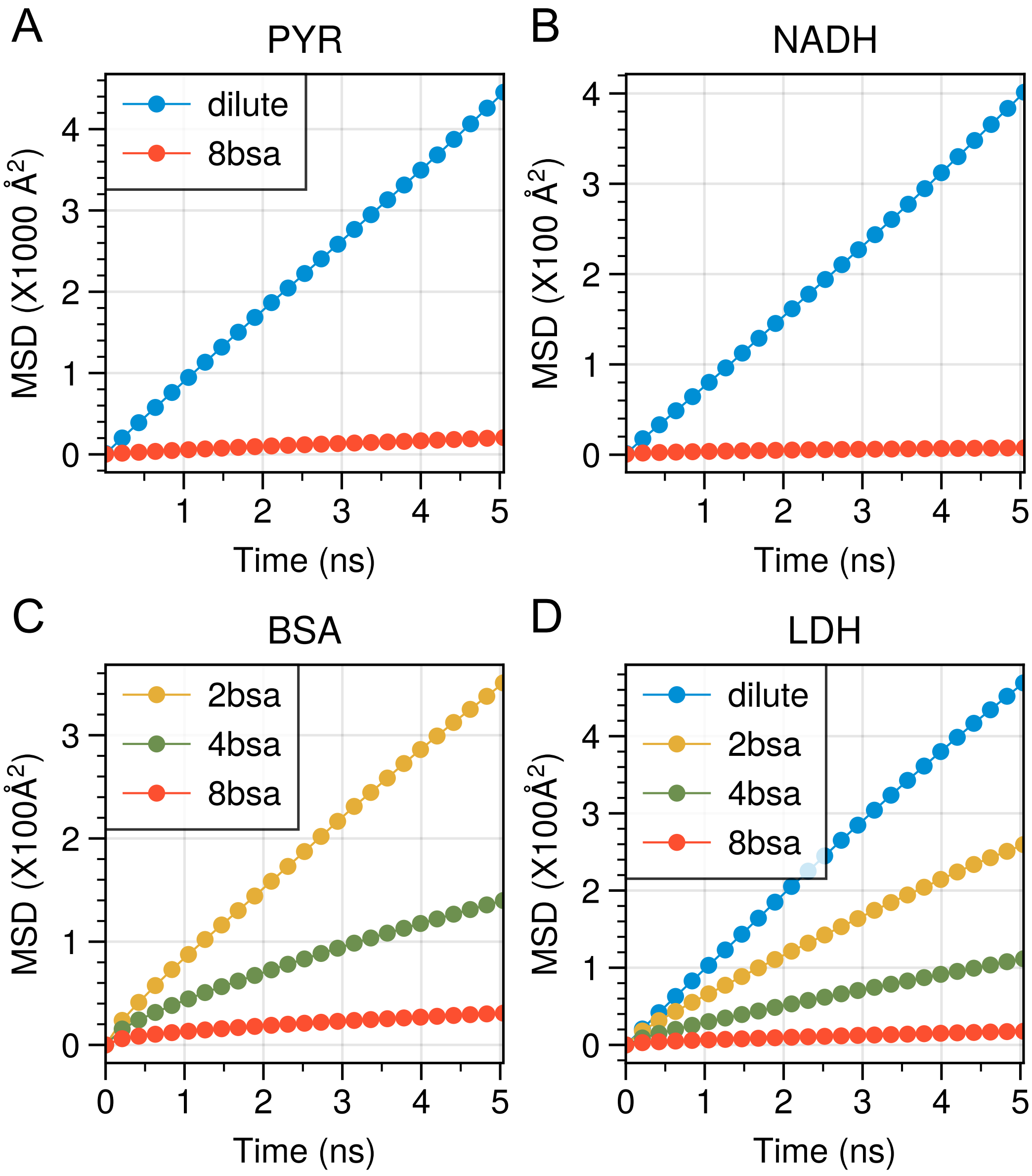


FIG. S9. **The mean squared displacement (MSD) of substrates (pyruvate and NADH), LDH tetramer, and crowder agent (BSA) in dilute and crowded conditions.** The crowded systems with 2, 4, 6, and 8 BSA molecules are denoted 2bsa, 4bsa, 6bsa, and 8bsa, respectively.

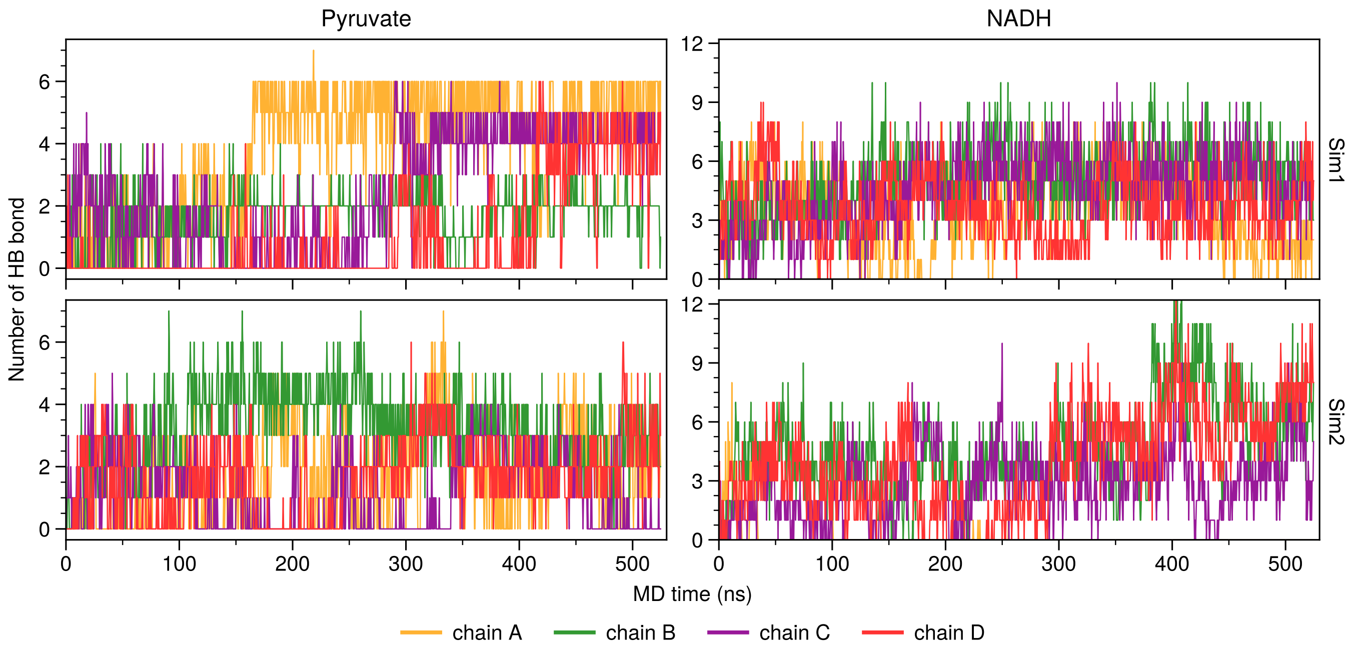


FIG. S10. **The number of hydrogen bonds formed between substrates (pyruvate and NADH) and the crowder agent (BSA) under crowded environmental conditions.** The substrates of the four monomers are distinguished by different colors, and the results from the two independent simulations are presented separately.

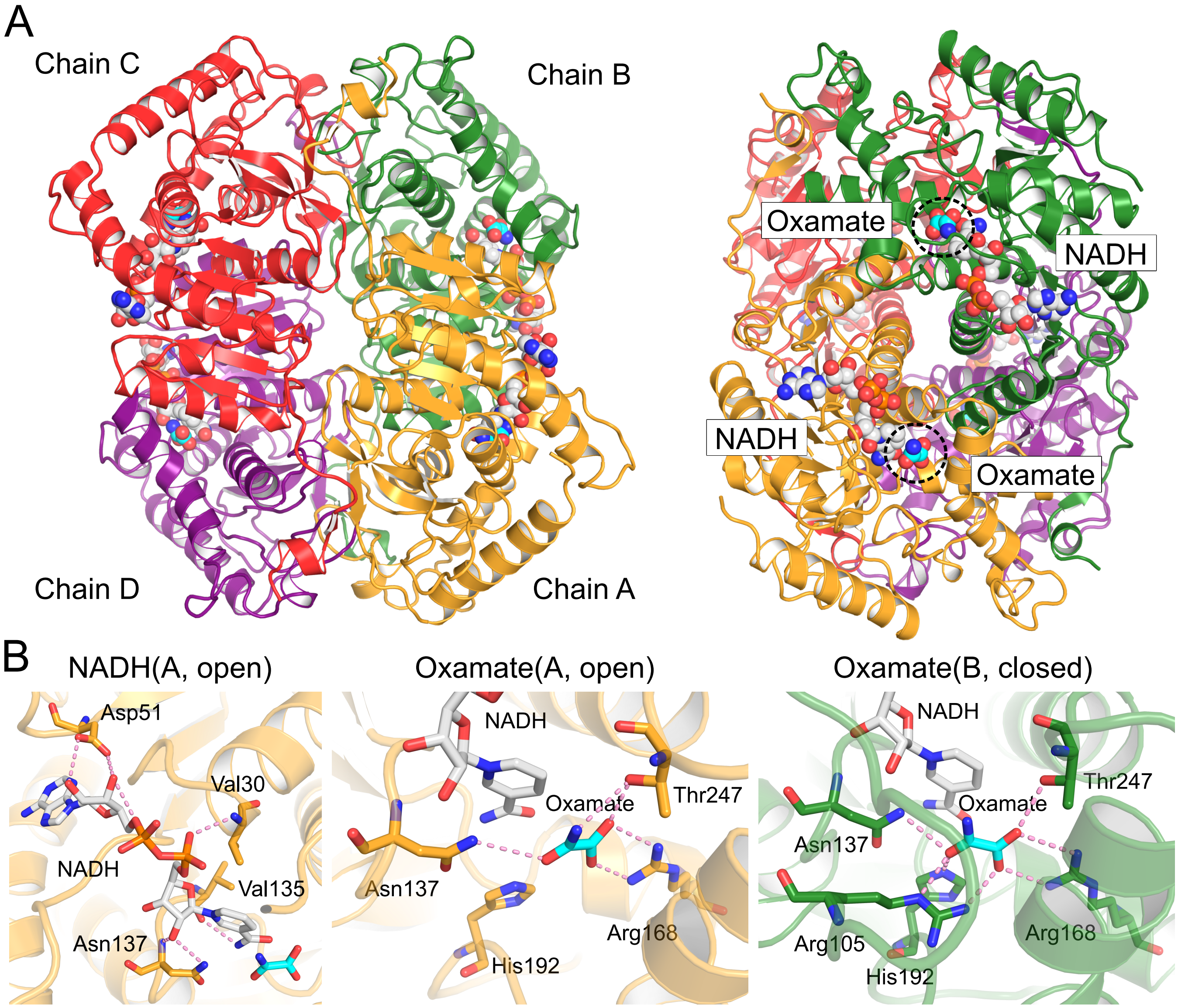


FIG. S11. **The crystal structure of rabbit muscle LDH tetramer (PDB code: 3H3F).** (A) Overview of the whole structure depicted from top (left) and side (right) with each monomer in different colors. NADH and oxamate (analogue of pyruvate) are illustrated with spheres. (B) Significant hydrogen-bond interactions between substrates and residues of LDH in open and closed forms.

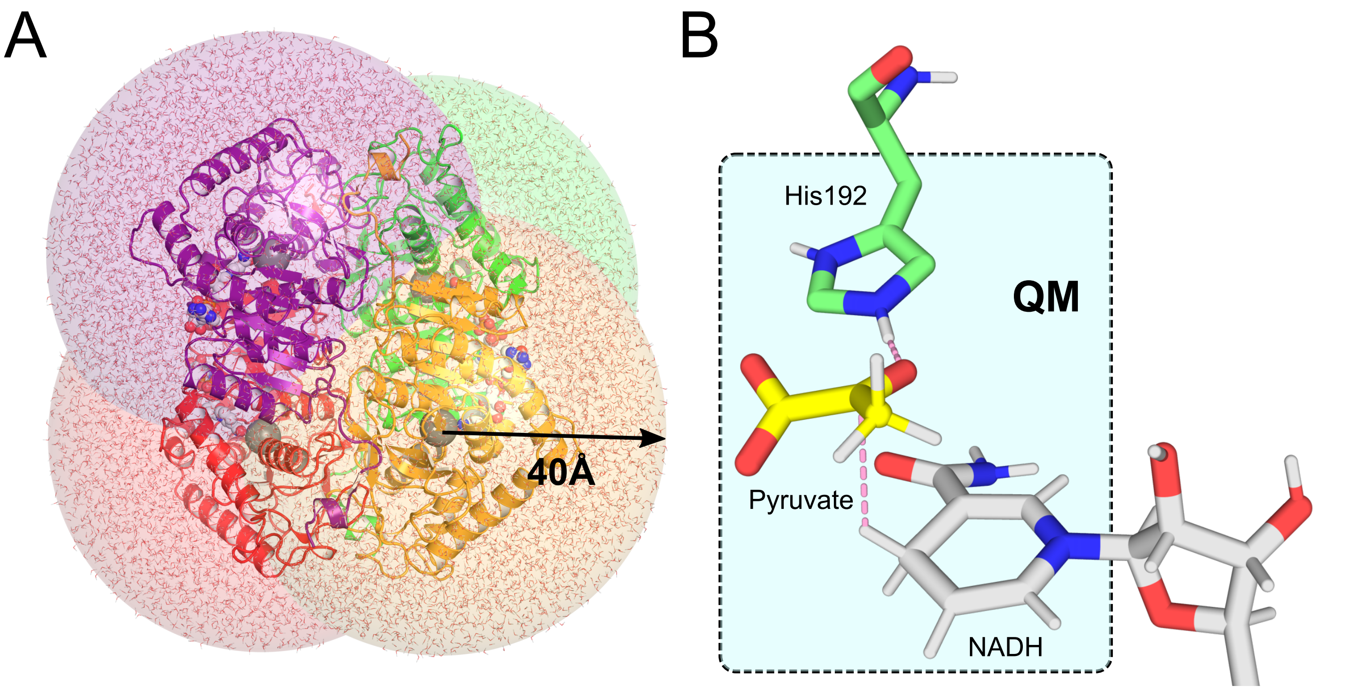


FIG. S12. **The QM/MM system used in the present study.** (A) The whole system is composed of four spheres with a radius of 40 Å, centered at the $C_\alpha$ atom of Ser160 of each LDH subunit. (B) The QM region is composed of pyruvate, the sidechain of His192, and a nicotinamide ring of NADH.

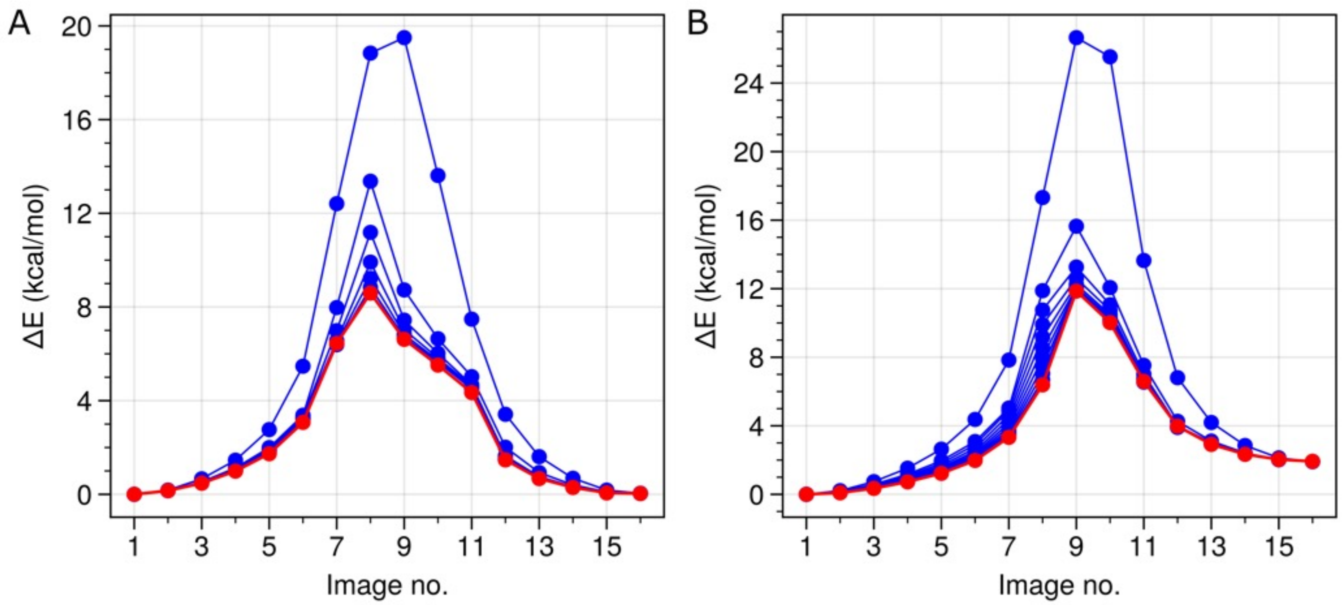


FIG. S13. **The convergence of the string simulations of the chemical reaction.** (A) Closed form and (B) open forms. The final pathway obtained from the string simulation is depicted in red.

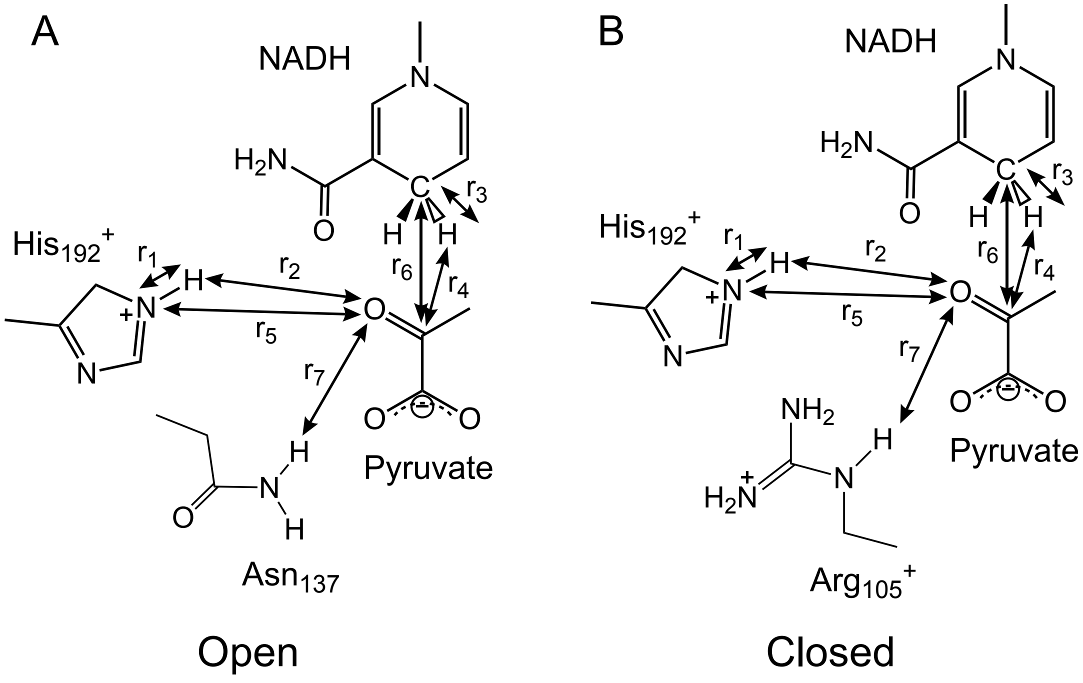


FIG. S14. **Illustration of collective variables (CVs) used for umbrella sampling of the chemical reaction.** The open (left) and closed(right) forms of LDH monomers.

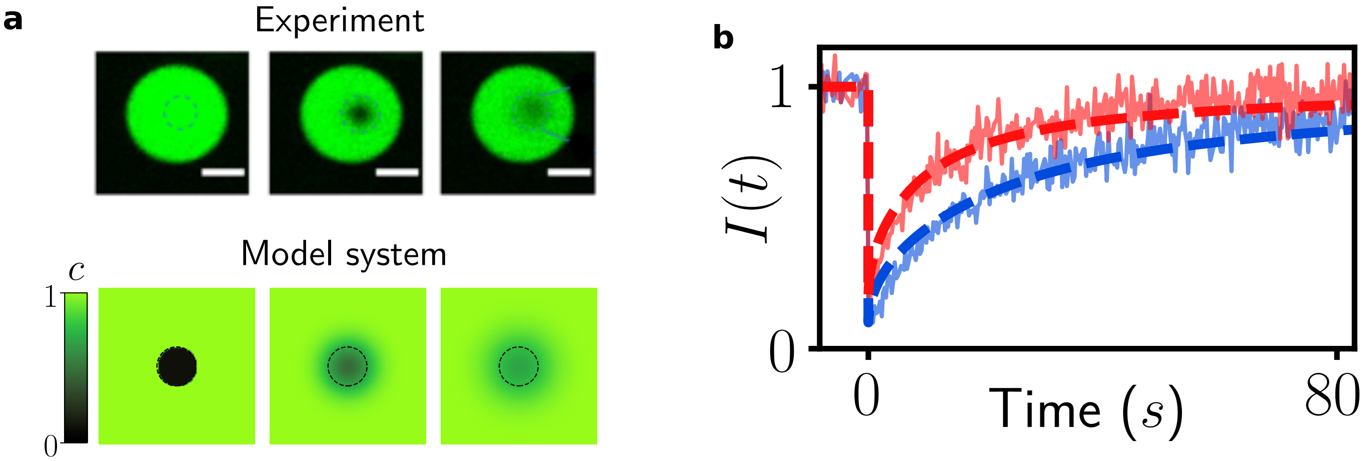


FIG. S15. **Results from the FRAP model system. a**, Top: images of fluorescent BSA in the experimental system during a bleach measurement. Bottom: concentration field during a simulated bleach in the theoretical model FRAP system described in Section II A. **b**, Intensity recovery curves from the model system (dashed lines) plotted on top of experimental data. Further discussion can be found in Section II B.

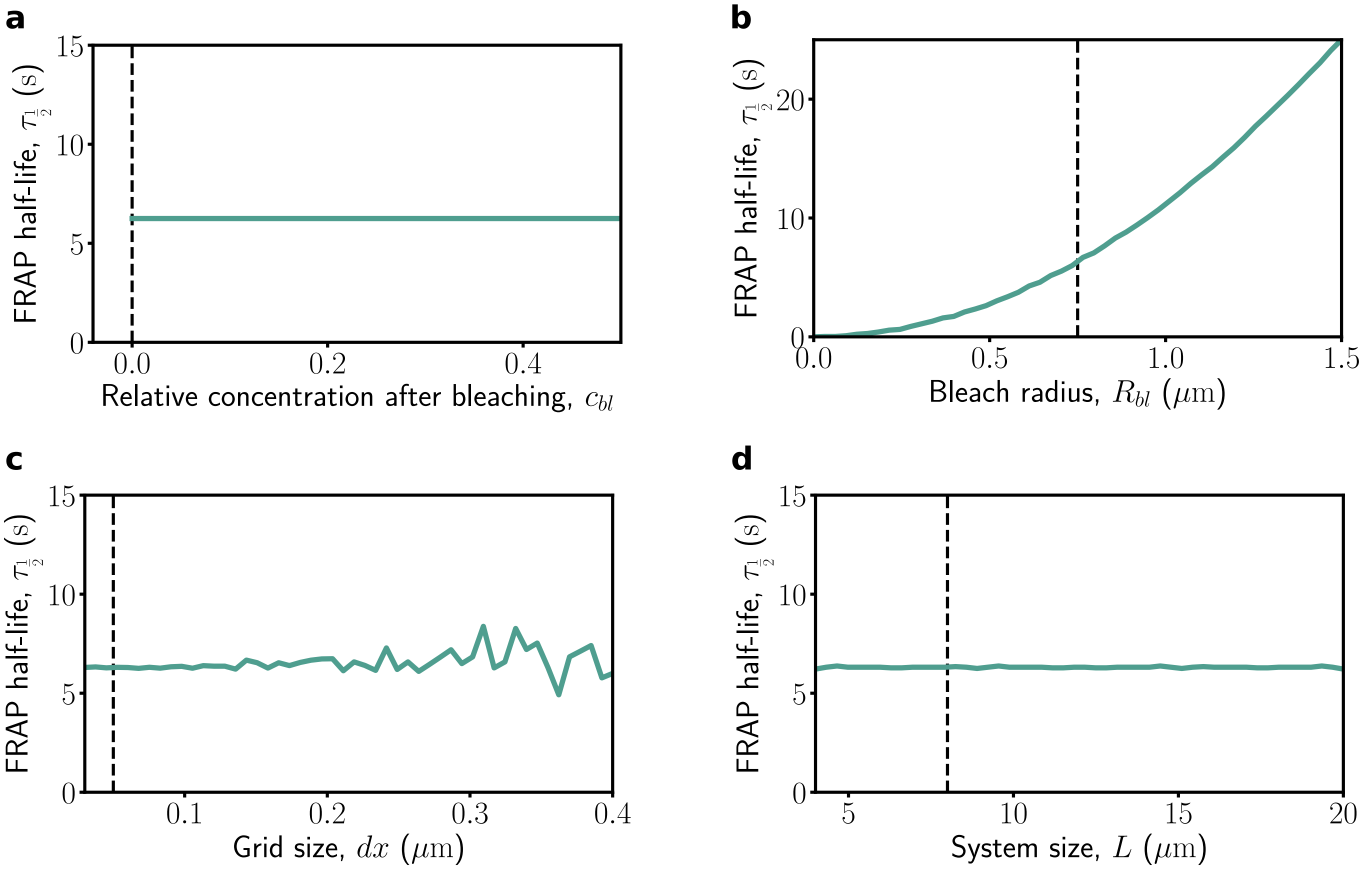


FIG. S16. **Validating the FRAP model system.** FRAP half-lives measured by numerically integrating the theoretical model system described in Section II A. The dashed vertical lines show the parameter values we use in the main text. Further discussion can be found in Section II B.

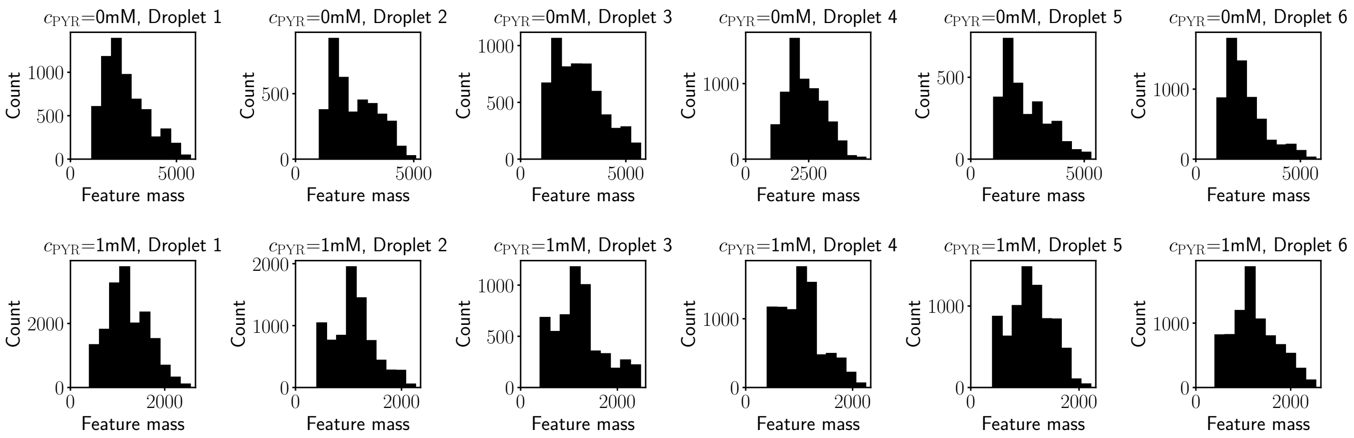


FIG. S17. **Feature mass distributions.** Distributions of the mass of detected features after filtering. As discussed in Section III A, we filter out any features with a brightness lower than 1000.

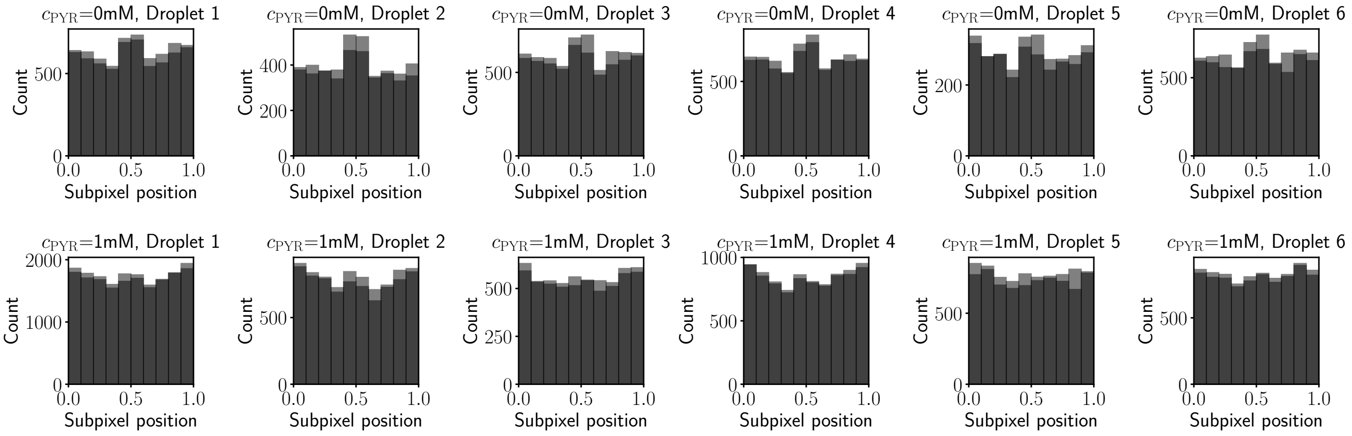


FIG. S18. **Subpixel position distributions.** Distributions of the subpixel positions of detected features after filtering. As discussed in Section III A, roughly uniform distributions are desired.

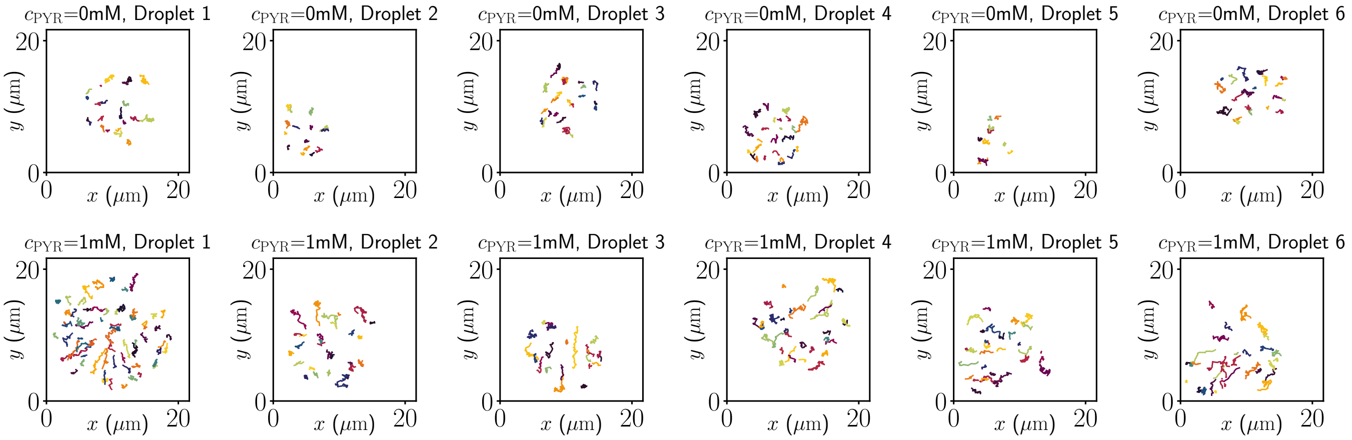


FIG. S19. **Extracted trajectories.** Trajectories extracted from inactive (0 mM pyruvate) and active (1 mM pyruvate) droplets, as described in Section III.

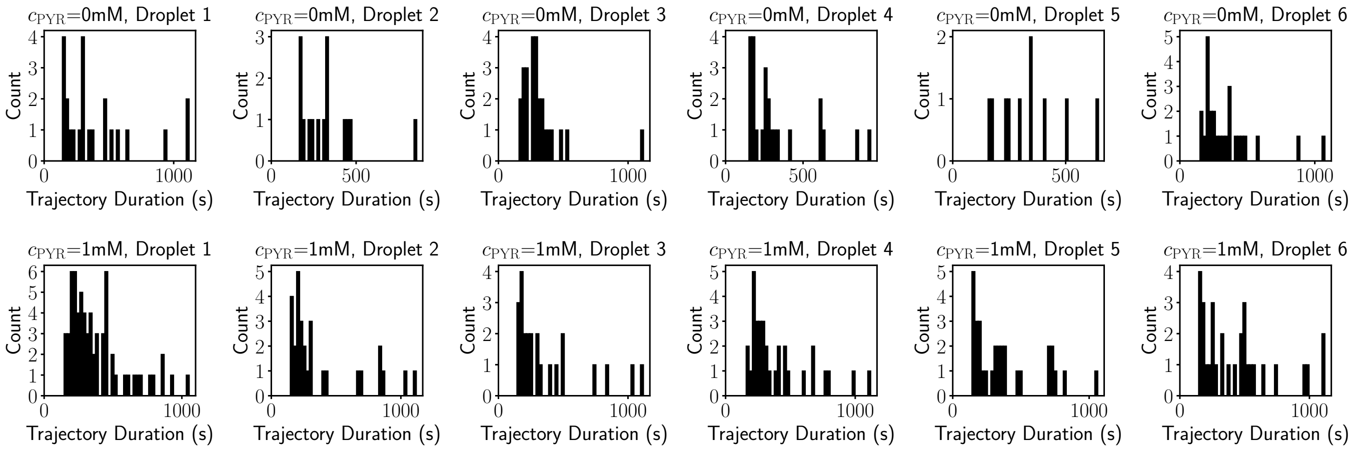


FIG. S20. **Trajectory duration distributions.** Distribution of the duration of the extracted trajectories. As discussed in Section III B we filter out any trajectories lasting less than 140 s.

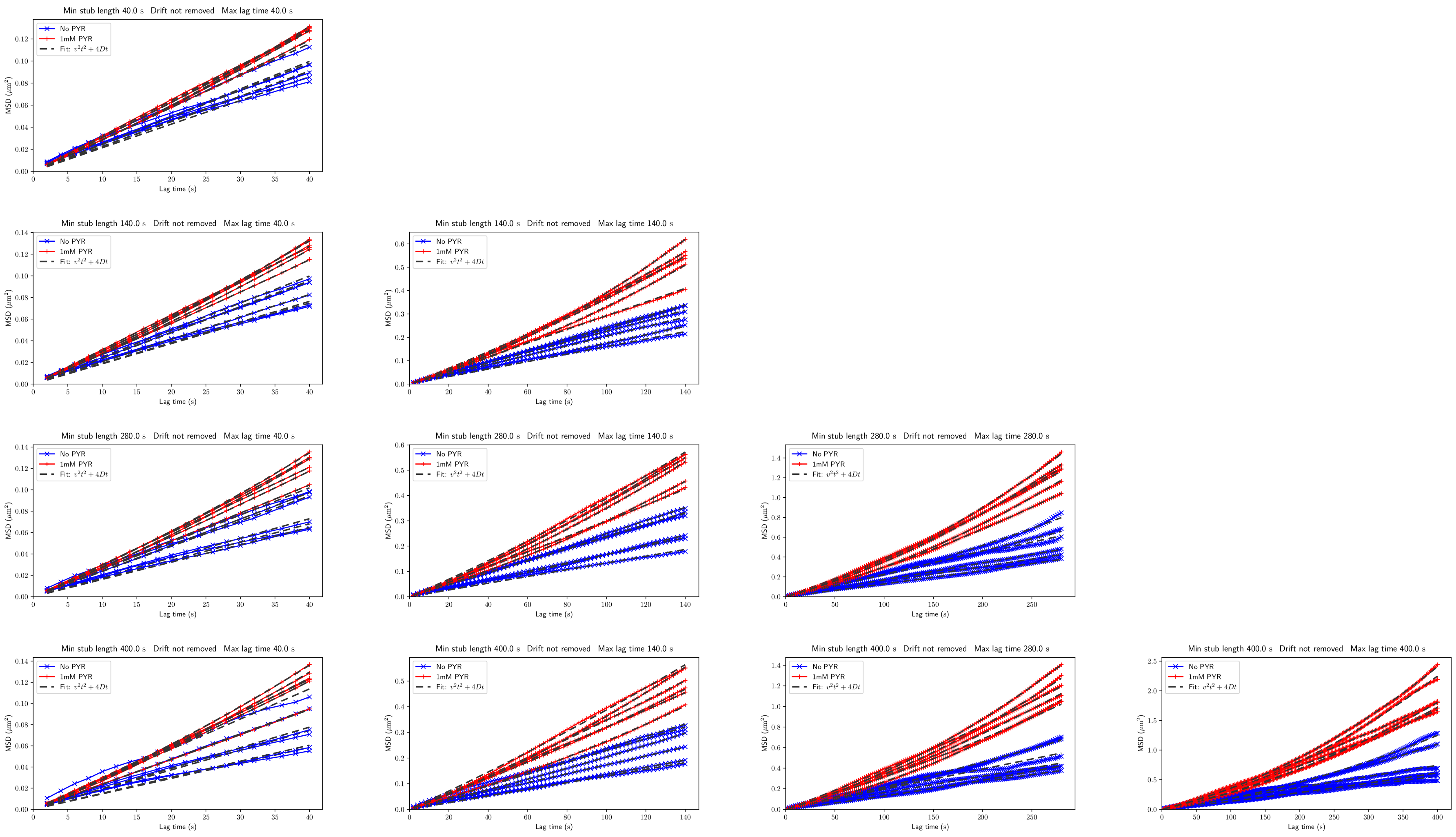


FIG. S21. **Validating the MSDs from nanoparticle tracking.** MSDs with different `min_stub_length` and `max_lag_time` applied. Further discussion can be found in Section III B. See Fig. S22 for the corresponding extracted parameters.

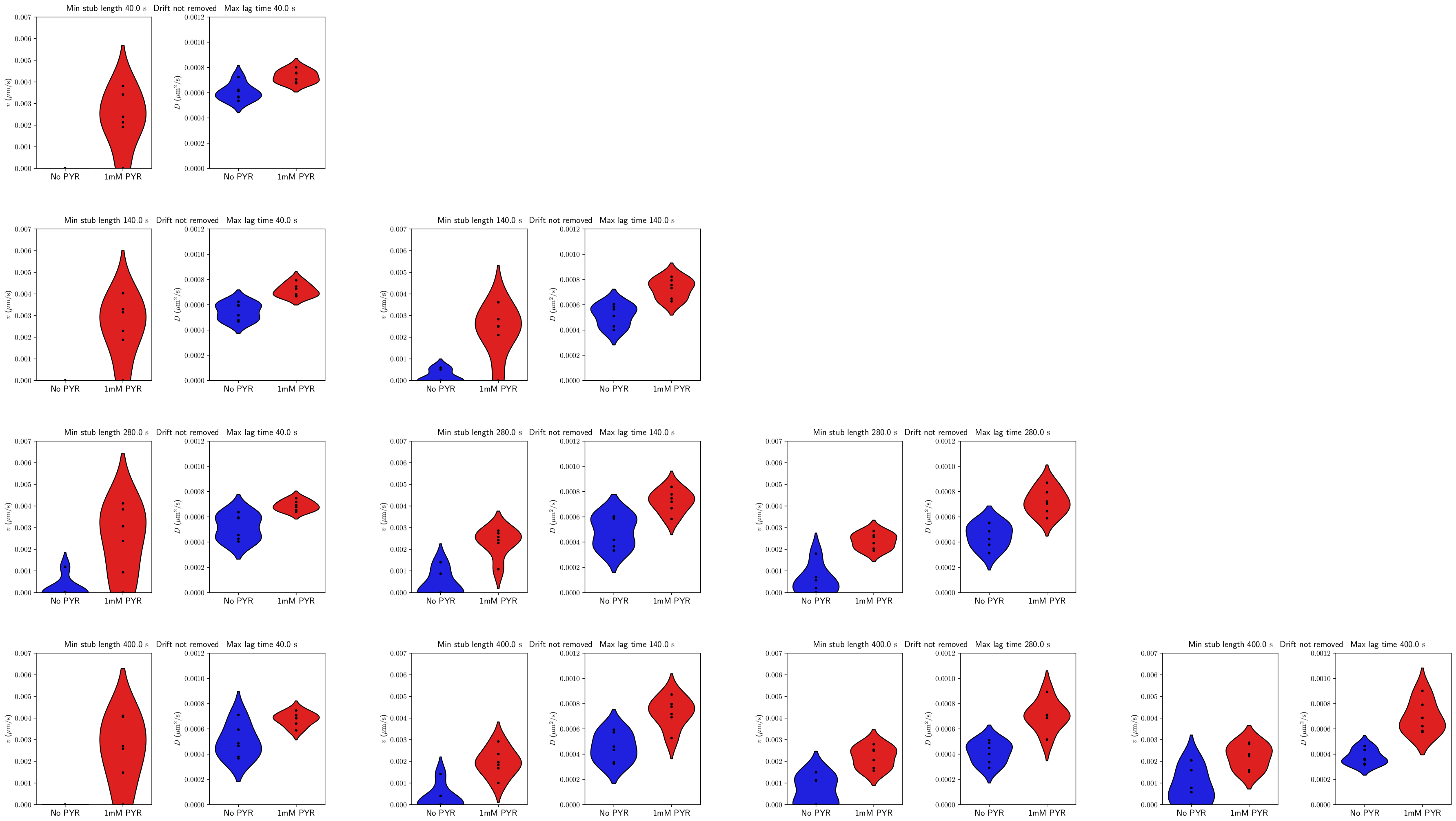


FIG. S22. **Validating the extracted parameters from nanoparticle tracking.** Extracted parameters with different `min_stub_length` and `max_lag_time` applied. Further discussion can be found in Section III B. See Fig. S21 for the corresponding MSDs.

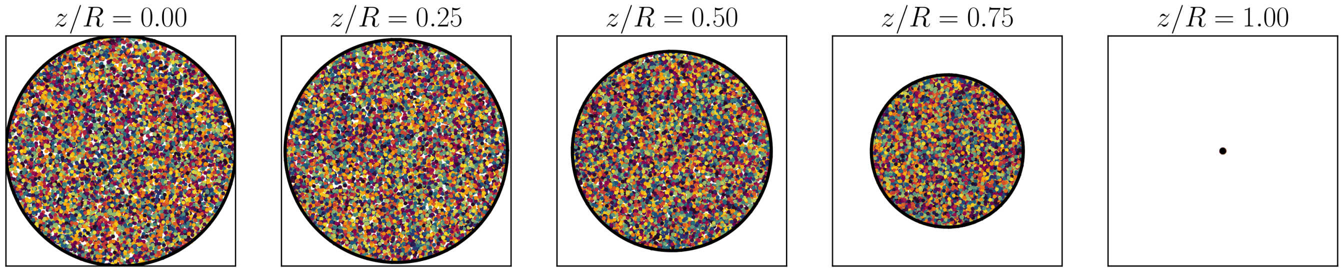


FIG. S23. **Monte Carlo sampling points.** Random sampling points used for the Monte Carlo calculation of projected velocity in Section IV C, shown for five different $z$ slices. We use $10^2$ slices with $10^5$ samples per slice.

#### II. FRAP ANALYSIS

We construct a model FRAP system to investigate the effect of changing $D$ and $v$ on the FRAP results. We then apply the FRAP model to extract the diffusion coefficient of BSA in active and inactive droplets.

##### A. Theoretical FRAP model

To model the FRAP experiments, we consider a two-dimensional system where fluorescent particles are diffusing subject to a drift of speed $v$. The concentration of fluorescent particles $c(\mathbf{r}, t)$ obeys the drift-diffusion equation

$$\partial_t c + \mathbf{v} \cdot \nabla c - D\nabla^2 c = 0. \tag{S1}$$

Although our calculation predicts a non-uniform flow profile, we can treat the flow in the area around the bleached region as approximately uniform in space. Therefore we take $\mathbf{v} = v\hat{\mathbf{v}}$ to be uniform in space, as well as constant in time. We assume an instantaneous bleach at $t = 0$, with a uniform bleached region of radius $R_{\mathrm{bl}}$ centred on the origin. We also assume that at $t = 0$ the concentration of fluorescent particles falls uniformly to $c_{\mathrm{bl}}$ inside the bleached region and that the concentration of fluorescent particles outside the bleached region is initially constant. This leads to the following initial condition

$$c(\mathbf{r}, t = 0) = \begin{cases} c_{\mathrm{bl}}, & r \leq R_{\mathrm{bl}} \\ c_0, & r > R_{\mathrm{bl}}. \end{cases} \tag{S2}$$

We assume that the intensity measured over a region is proportional to the concentration of fluorescent particles in that region, such that the normalised intensity over the initially bleached region $\mathscr{B}$ is

$$I_B(t) = \frac{\iint_{\mathscr{B}} c(\mathbf{r}, t)\ \mathrm{d}^2\mathbf{r}}{\iint_{\mathscr{B}} c_0\ \mathrm{d}^2\mathbf{r}}. \tag{S3}$$

We define the half-life $\tau_{\frac{1}{2}}$ as the time taken for the intensity over the initially bleached region to reach half the difference between its initial unbleached and bleached values.

$$I_B(\tau_{\frac{1}{2}}) = \frac{1}{2} \frac{\iint_{\mathscr{B}} c_0\ \mathrm{d}^2\mathbf{r} - \iint_{\mathscr{B}} c_{\mathrm{bl}}\ \mathrm{d}^2\mathbf{r}}{\iint_{\mathscr{B}} c_0\ \mathrm{d}^2\mathbf{r}} \tag{S4}$$

$$= \frac{1}{2}\left(1 - \frac{c_{\mathrm{bl}}}{c_0}\right). \tag{S5}$$

##### B. Numerical integration

This system can be numerically integrated very efficiently using vectorization. We use forward-difference time derivatives and central-difference spatial derivatives to calculate $c(\mathbf{r}, t)$.

In Fig. S15**a** we show the concentration field results during a simulated bleach in the model FRAP system alongside experimental images. The model faithfully captures the dynamics seen in experiments. From the concentration field we can then extract the intensity curve $I_B$, as described in the previous section. Recovery curves from the model are plotted on top of data from experiments in Fig. S15**b**, again showing how the model accurately reflects the experimental measurements.

From the intensity curves, for each set of parameters we can calculate the half-life. In the main text we investigate relationship between $D$ and $v$ on the half-lives, for example Fig. 3**b** of the main text. Here we verify the other system parameters that we use for the model system. In Fig. S16 we show the dependence of half-life on different system parameters. Due to the normalisation and definition of half-life, the half-lives do not depend on $c_{\mathrm{bl}}$, the concentration that the bleached region is bleached to. This is demonstrated in Fig. S16**a**. As shown in Fig. S16**b**, the radius of the bleached region $R_{\mathrm{bl}}$ does affect the half-life, so we use the same radius $R_{\mathrm{bl}} = 0.75\,\mu\mathrm{m}$ used in the FRAP experiments. Using a grid size $dx$ that is too large leads to instability in the numerical integration, which can be seen in Fig. S16**c**. Therefore, we ensure that the grid size we use $dx = 0.05\,\mu\mathrm{m}$ is much smaller than the threshold for stable results. The amount of bulk surrounding the bleached region is represented by the system size $L$. Although we use a value $L = 8\,\mu\mathrm{m}$ to reflect the approximate size of the droplets, as shown in Fig. S16**d** the value of $L$ does not affect the half-lives. The values of each of these system parameters that we use in the main text are marked by the dashed lines in Fig. S16.

### III. NANOPARTICLE TRACKING

We analyse videos of nanoparticle tracers to extract their diffusion constant and drift speed in active and inactive droplets.

#### A. Obtaining trajectories

This first step in the tracking analysis is to crop out an individual droplet from the full video. We took droplets which were as isolated as possible, as the FRAP measurements were made on isolated droplets. Furthermore we excluded any droplets showing significant anomalous movement of the entire droplet. We then use the `trackpy` Python package [1], later abbreviated to `tp`, to identify and track the nanoparticles in the videos. Cleaning the data and isolating the proper particle tracks requires a few steps and various parameters, which we detail here.

We load in the frames of the video, taking the intensity of the green channel to give a greyscale image. The videos are 0.5 fps, meaning each frame corresponds to 2 s. The pixels are square, with side length 0.22 µm.

There are two important parameters when locating particles, also referred to as features, in a frame: the features' `diameter` and the minimum integrated brightness of a feature `minmass`. We set `diameter` to be large enough to avoid subpixel bias. Plot of the subpixel position distributions are shown in Fig. S18. We set `minmass` to be above the large, low mass peak. This removes unhelpful small patches of brightness. The mass distribution of features following this filtering is shown in Fig. S17.

After using `tp.batch` to locate the features in the different frames, to obtain trajectories we need to link particle identities between frames, which is done using `tp.link`. We set `search_range`, the distance that particles can move between frames, to 2 px, which corresponds to a maximum speed of $0.2\,\mu\mathrm{m\,s^{-1}}$. Another parameter here is `memory`, which we set to 2. This means that particles can disappear and reappear up to 4 s later and still be counted as the same particle. We then use `tp.filter_stubs` to only consider trajectories which last at least as long as the maximum MSD lag-time.

The extracted trajectories are shown in Fig. S19. Videos of the extracted trajectories overlaid onto the original frames can be seen in the video file *Annotated_combined.mp4*. We show the distributions of trajectory durations for each droplet in Fig. S20. With the parameters we use we obtain a good number of long-lasting trajectories in each droplet.

#### B. Calculating MSDs and extracting parameters

Given the particle trajectories, we then use `tp.emsd` to calculate the ensemble mean-squared displacement. We fit the resulting MSDs to theoretical expression for the mean-squared displacement

$$\mathrm{MSD} = v^2 t^2 + 4Dt \tag{S6}$$

and extract the parameters $v$ and $D$, enforcing $v \geq 0$ and $D \geq 0$.

In Fig. S21 we show MSDs and in Fig. S22 the corresponding extracted parameters, with different `min_stub_length` and `max_lag_time` applied. As part of the filtering we remove any trajectories shorter than `min_stub_length` and `max_lag_time` corresponds to the maximum lag time in the MSDs. As described in the previous subsection, we ensure that `min_stub_length` $\geq$ `max_lag_time`. From the MSDs we see that filtering to keep only very long times ($\gtrsim 200$ s) we do not have enough trajectories to give good quality statistics. Therefore we set `min_stub_length` $=$ `max_lag_time` $=$ 140 s. Looking at the extracted parameters, the results are robust to changes in the parameters. There are some exceptions at longer times, however as discussed in these cases the trajectories are not sufficient to give reliable results.

### IV. DERIVATION OF THE INDUCED VELOCITY FLOW FIELD

#### A. Hydrodynamics

Here we derive the solution of the velocity flow inside and outside the enzymatic droplet. We consider a hemispherical droplet of radius $R$ on a solid substrate with the plane of the surface being the $x-y$ plane and the height of the droplet is towards the $z$-direction. We start from the incompressible Stokes equation

$$-\eta \nabla^2 \mathbf{v} = -\nabla p, \tag{S7}$$

$$\nabla \cdot \mathbf{v} = 0. \tag{S8}$$

where $\eta$ is the viscosity of the fluid, $\mathbf{v}$ the velocity (flow) field and $p$ is the pressure. The incompressibility condition implies that pressure satisfies Laplace equation. By taking into account the symmetries of the problem that leads us to use the spherical polar coordinates $(r, \theta, \phi)$, we choose an *ansatz* for the pressure field as follows

$$p(r,\theta) = \begin{cases} \dfrac{\eta}{R}\left[A_{\text{in}} + B_{\text{in}}\left(\dfrac{r}{R}\right)^2 P_2(\cos\theta)\right], & r \leq R, \\ \dfrac{\eta}{R}\left[A_{\text{out}}\left(\dfrac{R}{r}\right) + B_{\text{out}}\left(\dfrac{R}{r}\right)^3 P_2(\cos\theta)\right], & r \geq R. \end{cases} \tag{S9}$$

where $P_2(\cos\theta)$ is the second Legendre polynomial (with $\theta$ being the polar angle measured from the $z$-axis), and $A_{\text{in}}$, $A_{\text{out}}$, $B_{\text{in}}$, and $B_{\text{out}}$ are unknown coefficients. We define the following quantity

$$\chi(r,\theta) = \frac{1}{R}(\mathbf{r}\cdot\mathbf{v}). \tag{S10}$$

and expand it in Legendre polynomials as

$$\chi(r,\theta) = \sum_{\ell} f^{(\ell)}(r) P_\ell(\cos\theta). \tag{S11}$$

Assuming that there is no source/sink at the center of the droplet and no flow at infinity as boundary conditions, we solve the differential equations, to obtain

$$f_{\text{in}}^{(2)}(r) = \frac{B_{\text{in}}}{7}\left[\left(\frac{r}{R}\right)^4 - \left(\frac{r}{R}\right)^2\right] + \left(C + \frac{B_{\text{out}}}{2}\right)\left(\frac{r}{R}\right)^2, \tag{S12}$$

$$f_{\text{out}}^{(2)}(r) = \frac{B_{\text{out}}}{2}\left(\frac{R}{r}\right) + C\left(\frac{R}{r}\right)^3, \tag{S13}$$

where $C$ is an unknown integration constant, and $A_{\text{in}} = A_{\text{out}} = 0$.

The above solution allows us to obtain the radial component of the velocity $v_r = \hat{\mathbf{r}}\cdot\mathbf{v}$. By utilizing the expressions for the stream function $\psi$ in axially-symmetric flows in three-dimensions [2]

$$v_r = -\frac{1}{r^2\sin\theta}\partial_\theta\psi, \tag{S14}$$

$$v_\theta = \frac{1}{r\sin\theta}\partial_r\psi, \tag{S15}$$

we find

$$v_r(r,\theta) = P_2(\cos\theta)\begin{cases} \dfrac{B_{\text{out}}}{4}\left[5\left(\dfrac{r}{R}\right) - 3\left(\dfrac{r}{R}\right)^3\right] + \dfrac{C}{2}\left[7\left(\dfrac{r}{R}\right) - 5\left(\dfrac{r}{R}\right)^3\right], & r \leq R, \\ \dfrac{B_{\text{out}}}{2}\left(\dfrac{R}{r}\right)^2 + C\left(\dfrac{R}{r}\right)^4, & r \geq R, \end{cases} \tag{S16}$$

$$v_\theta(r,\theta) = \frac{P_3(\cos\theta) - P_1(\cos\theta)}{5\sin\theta}\begin{cases} \dfrac{15B_{\text{out}}}{4}\left[\left(\dfrac{r}{R}\right) - \left(\dfrac{r}{R}\right)^3\right] + \dfrac{C}{2}\left[21\left(\dfrac{r}{R}\right) - 25\left(\dfrac{r}{R}\right)^3\right], & r \leq R, \\ -2C\left(\dfrac{R}{r}\right)^4, & r \geq R, \end{cases} \tag{S17}$$

where we have eliminated $B_{\text{in}}$ by using continuity of $v_\theta$ at $r = R$ as a boundary condition.

##### B. Connecting flow velocity and concentration

For the concentration, we consider a scalar field $c(\mathbf{r}, t)$ that satisfies a reaction-diffusion equation given by,

$$\partial_t c - D\nabla^2 c = \begin{cases} k_{\text{cat}}\rho, & r \leq R \\ 0, & r \geq R, \end{cases} \tag{S18}$$

where $D = \frac{k_B T}{6\pi\eta\ell}$ is the diffusion coefficient of the chemical molecules that have a hydrodynamic radius $\ell$, $k_{\text{cat}}$ is the catalytic reaction rate of the enzymes, and $\rho$ is the density of the enzymes in the droplet. In steady-state, the above equation reduces to a Poisson equation. Using the symmetries of the problem, we choose the following form for the enzyme density

$$\rho(r,\theta) = \begin{cases} \rho_m \left[ \left(1 - \left(\frac{r}{R}\right)^2\right) + b\cos^2\theta \left(\frac{r}{R}\right)^2 \right], & r \le R, \\ 0, & r \ge R, \end{cases} \tag{S19}$$

where $b$ is a positive constant to be determined via consistency conditions. The form of the enzyme density is justified since a higher enzyme density is expected at the centre of the droplet, while the symmetries of the problem allow for a lowest harmonic axisymmetric variation in the enzyme density. The coefficients are chosen such that a self-consistent exact solution to the complete problem can be obtained.

The presence of the surface substrate in combination with the enzymatic activity induces a phoretic slip velocity $v_s$ near the surface, which is proportional to gradient of the concentration $c$. Hence, by solving Eq. (S19) we connect the emerging phoretic velocity with our hydrodynamic solution through the boundary condition

$$v_s \equiv v_r(r, \theta = \pi/2) = \mu \partial_r c(r). \tag{S20}$$

where the phoretic mobility is given as $\mu = \frac{k_B T}{\eta}\lambda_D^2$ in terms of the Derjaguin length $\lambda_D$, which measures the range of the intermolecular interaction between the chemical and the surface [3].

Using the surface slip-velocity boundary condition, we can find the solution for the radial and polar velocity components as

$$v_r(r,\theta) = v_0 \left(3\cos^2\theta - 1\right) \begin{cases} \left[ \left(\frac{1}{3} + \frac{b}{15}\right)\left(\frac{r}{R}\right) - \left(\frac{1}{5} + \frac{b}{35}\right)\left(\frac{r}{R}\right)^3 \right], & r \le R, \\ \left[ \left(\frac{2}{15} + \frac{b}{15}\right)\left(\frac{R}{r}\right)^2 - \left(\frac{b}{35}\right)\left(\frac{R}{r}\right)^4 \right], & r \ge R, \end{cases} \tag{S21}$$

and

$$v_\theta(r,\theta) = v_0 \left(-\sin\theta\cos\theta\right) \begin{cases} \left[ \left(1 + \frac{b}{5}\right)\left(\frac{r}{R}\right) - \left(1 + \frac{b}{7}\right)\left(\frac{r}{R}\right)^3 \right], & r \le R, \\ \frac{2b}{35}\left(\frac{R}{r}\right)^4, & r \ge R, \end{cases} \tag{S22}$$

where the velocity scale is given as

$$v_0 = \frac{\mu k_{\text{cat}} \rho_m R}{D} = 6\pi k_{\text{cat}} \ell \rho_m \lambda_D^2 R. \tag{S23}$$

Using typical values for the parameters $\lambda_D \sim 1\,\text{nm}$, $\ell \sim 1\,\text{nm}$, $R \sim 10\,\mu\text{m}$, $k_{\text{cat}} \sim 10^3\ \text{s}^{-1}$ and $\rho_{\text{m}} \sim 1\,\mu\text{M}$, we find the value of $v_0 \simeq 0.1\,\mu\text{m}\,\text{s}^{-1}$.

Tangential stress balance at the boundary can be achieved via the gradient of the surface tension $\gamma$, which will depend on the local value of the concentration, leading to the following condition

$$\sigma_{r\theta}^{\text{in}} - \sigma_{r\theta}^{\text{out}} = -\frac{1}{R}\frac{\partial\gamma}{\partial c}\partial_\theta c, \tag{S24}$$

where,

$$c(r,\theta) = \frac{k_{\text{cat}}\rho_m R^2}{D}\left[ \left(\frac{1}{4} + \frac{b}{12}\right) - \left(\frac{1}{6} + \frac{b}{30}\right)\left(\frac{r}{R}\right)^2 + \left(\frac{1}{20} + \frac{b}{140}\right)\left(\frac{r}{R}\right)^4 + b\cos^2\theta\left(\frac{1}{10}\left(\frac{r}{R}\right)^2 - \frac{1}{14}\left(\frac{r}{R}\right)^4\right)\right] \tag{S25}$$

and

$$\sigma_{r\theta}(r,\theta) = \frac{k_B T \lambda_D^2 k_{\text{cat}} \rho_m}{D}(-\cos\theta\sin\theta) \begin{cases} \left[ \left(\frac{10+2b}{5}\right) - \left(\frac{112+16b}{35}\right)\left(\frac{r}{R}\right)^2 \right], & r \le R \\ \left(\frac{4+2b}{5}\right)\left(\frac{R}{r}\right)^3 - \frac{16b}{35}\left(\frac{R}{r}\right)^5, & r \ge R. \end{cases} \tag{S26}$$

Therefore, the expression for $b$ is found as

$$b = 35 \cdot \frac{k_\mathrm{B} T \lambda_\mathrm{D}^2}{R} \left(\frac{\partial \gamma}{\partial c}\right)^{-1} . \tag{S27}$$

Using $k_\mathrm{B}T = 4 \times 10^{-21}\,\mathrm{kg\,m^2\,s^{-2}}$, $\lambda_\mathrm{D} = 1\,\mathrm{nm}$, $R = 10\,\mathrm{\mu m}$, and $\partial\gamma/\partial c \simeq 2 \cdot 10^{-23}\,\mathrm{kg\,m^3\,s^{-2}}$ [4], we find

$$b \simeq 10^{-8}, \tag{S28}$$

which means that it can be safely neglected in the above results for practical purposes. This approximation leads to the results reported in the main text.

#### C. Projected velocity

The fluorescence images show the motion of the tagged particles in the horizontal plane. Therefore, we project our solution to obtain the projected flow velocity $v$ in the $x - y$ plane. The amplitude of the projected velocity is given by the following expression

$$v(\rho, z) = |v_r(r,\theta)\sin\theta + v_\theta(r,\theta)\cos\theta| = \left| v_r(\rho, z)\frac{\rho}{\sqrt{\rho^2+z^2}} + v_\theta(\rho, z)\frac{z}{\sqrt{\rho^2+z^2}} \right|, \tag{S29}$$

where $z = r\cos\theta$ and $\rho = r\sin\theta$. Using this expression, we can calculate the average velocity at different heights in the droplet upon integration, as follows

$$\sqrt{\langle v^2 \rangle}(z) = \left[ \frac{1}{\pi(R^2 - z^2)} \int_0^{2\pi} \mathrm{d}\phi \int_0^{\sqrt{R^2-z^2}} \mathrm{d}\rho \left(v^2\,\rho\right) \right]^{\frac{1}{2}} . \tag{S30}$$

We confirm our calculation using Monte Carlo random sampling algorithm, as shown in Fig. 3**i** in the main text. The sampling points used for the Monte Carlo calculation are shown at five different $z$ slices in Fig. S23.

---


[1] D. B. Allan, T. Caswell, N. C. Keim, C. M. van der Wel, and R. W. Verweij, soft-matter/trackpy: v0.6.4 (2024).
[2] J. Happel and H. Brenner, *Low Reynolds Number Hydrodynamics* (Springer Netherlands, 1983).
[3] R. Golestanian, Phoretic Active Matter, in *Active Matter and Nonequilibrium Statistical Physics: Lecture Notes of the Les Houches Summer School: Volume 112, September 2018* (Oxford University Press, 2022).
[4] A. Testa, M. Dindo, A. A. Rebane, B. Nasouri, R. W. Style, R. Golestanian, E. R. Dufresne, and P. Laurino, Sustained enzymatic activity and flow in crowded protein droplets, Nat. Commun. **12**, 1 (2021).
