## Extended Figures for "Enzymes can activate and mobilize the cytoplasmic environment across scales"

*Extended Data*

Mirco Dindo,[1,2] Jakob Metson,[3] Weitong Ren,[4] Michalis Chatzittofi,[3] Kiyoshi Yagi,[4] Yuji Sugita,[4,5,6,*] Ramin Golestanian,[3,7,†] and Paola Laurino[1,8,‡]

[1] *Protein Engineering and Evolution Unit, Okinawa Institute of Science and Technology Graduate University; Onna, Okinawa, Japan*
[2] *Department of Medicine and Surgery, Section of Physiology and Biochemistry, University of Perugia, Perugia, Italy*
[3] *Max Planck Institute for Dynamics and Self-Organization (MPI-DS), 37077 Göttingen, Germany*
[4] *Theoretical Molecular Science Laboratory, RIKEN Cluster for Pioneering Research, Saitama, Japan*
[5] *Computational Biophysics Research Team, RIKEN Center for Computational Science, Hyogo, Japan*
[6] *Laboratory for Biomolecular Function Simulation, RIKEN Center for Biosystems Dynamics Research, Hyogo, Japan*
[7] *Rudolf Peierls Centre for Theoretical Physics, University of Oxford, Oxford OX1 3PU, United Kingdom*
[8] *Institute for Protein Research, Osaka University, Suita 565-0871, Japan*

(Dated: January 28, 2025)

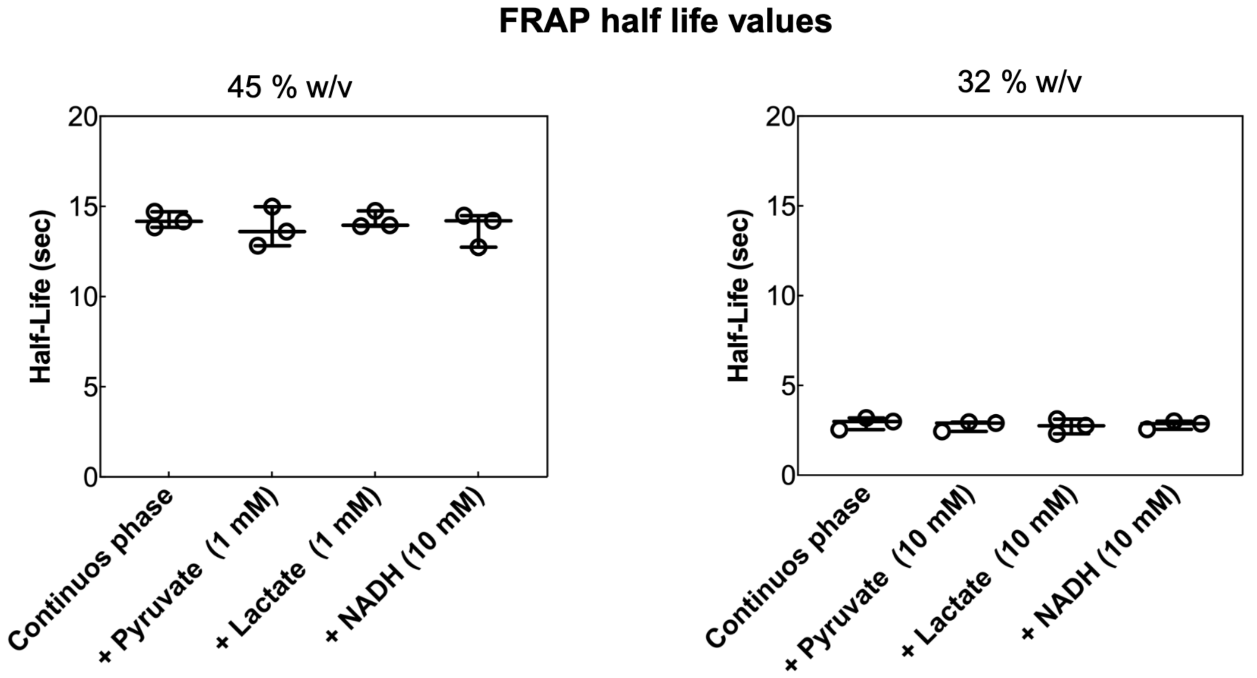


Extended Data Fig. E1. **Half-life values obtained by FRAP using a passive chemical gradient on the inactive droplets**. Plots of the half-life values (expressed in seconds) obtained from FRAP measurements on enzymatically inactive droplets in presence of several small molecules at different concentrations and containing different BSA content percentange (45% and 32% BSA (w/v)). The small molecules used during the FRAP experiments were pyruvate (1 and 10 mM, LDH reaction substrate), lactate (1 and 10 mM, LDH reaction product) and cofactor NADH (10 mM).

*
†
‡

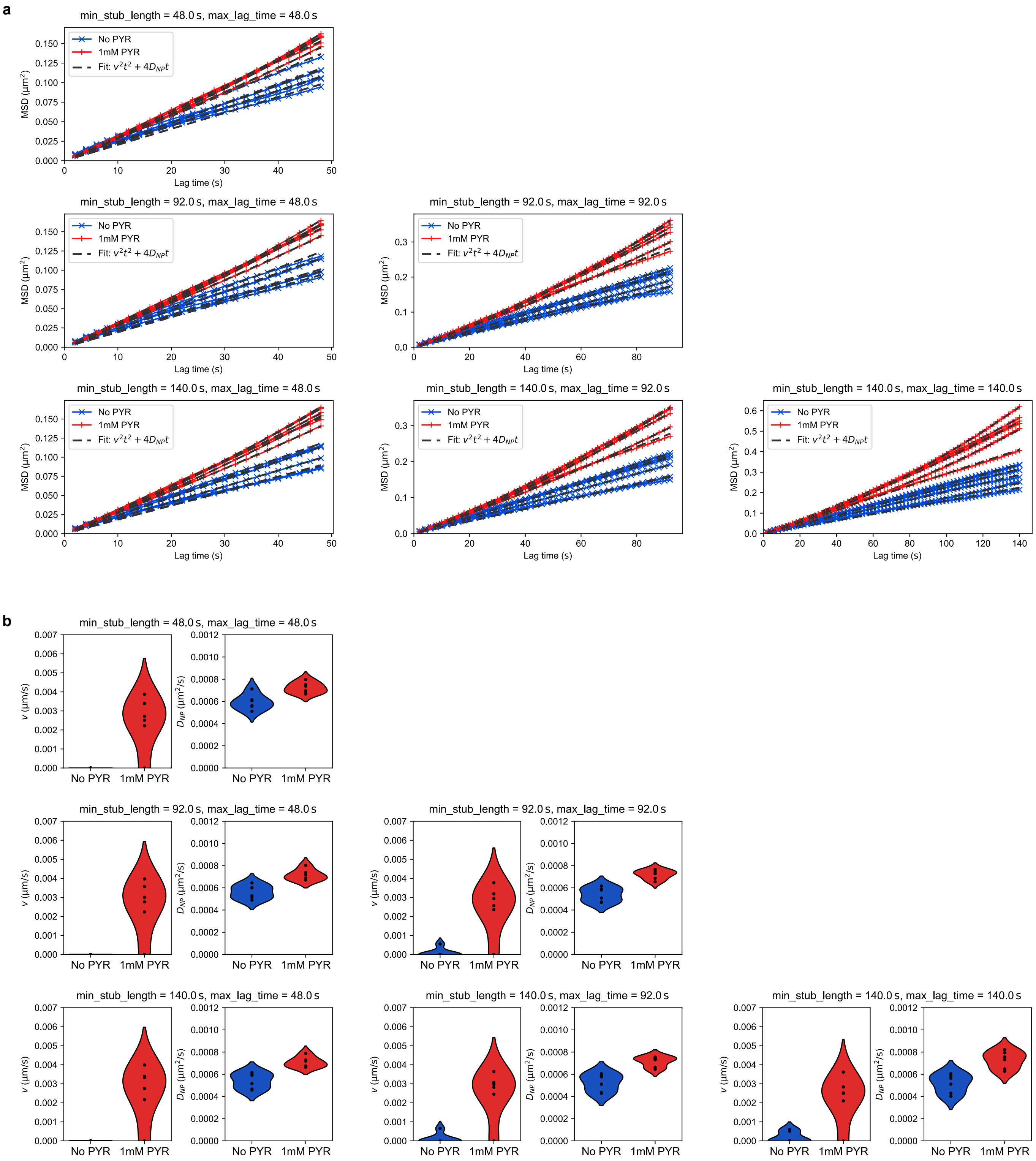


Extended Data Fig. E2. **Validating MSDs and extracted parameters from nanoparticle tracking.** **a**, MSDs and **b**, extracted parameters with different min_stub_lengths and max_lag_times applied. min_stub_length is the minimum length of retained trajectories and max_lag_time corresponds to the maximum lag time in the MSDs. The extracted parameters remain robust as min_stub_length and max_lag_time are varied. Further discussion can be found in the Supplementary Information.

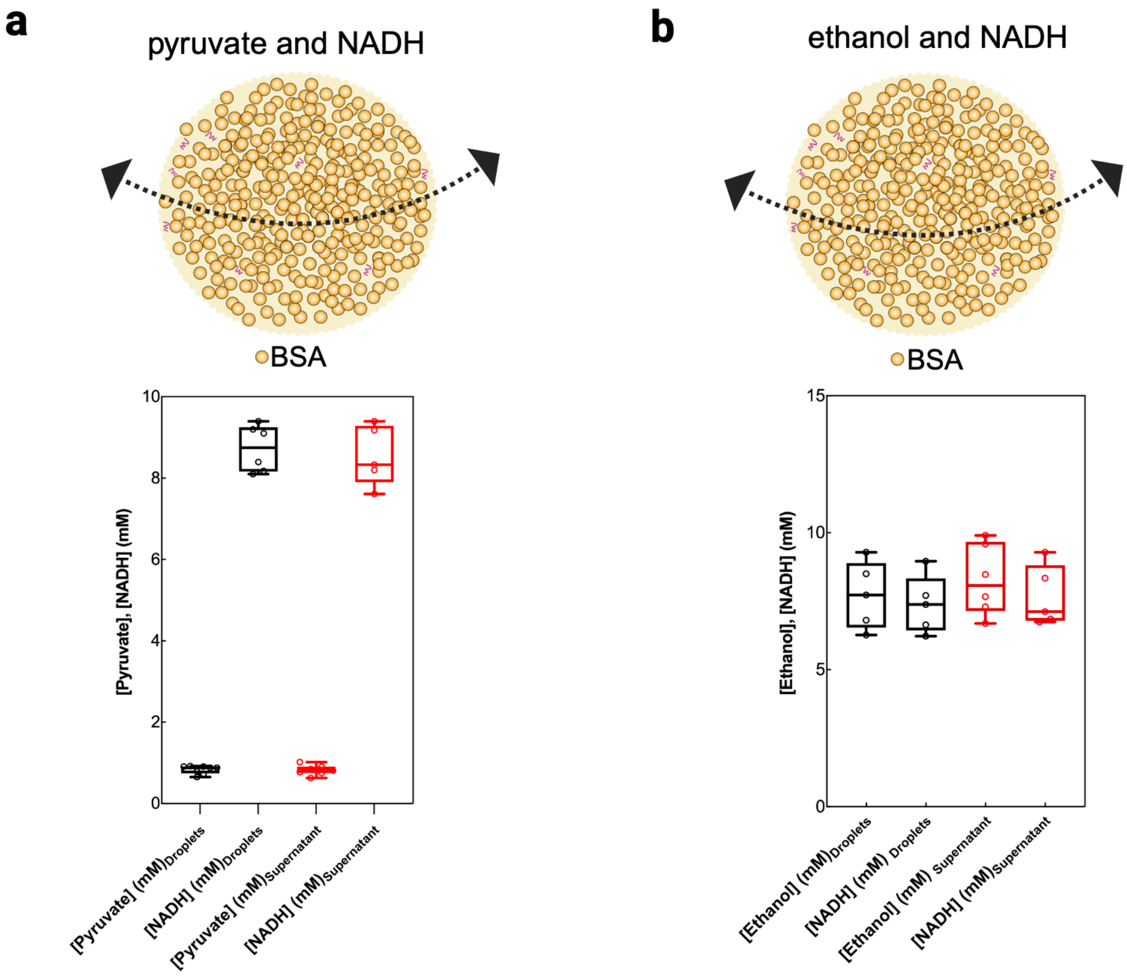


Extended Data Fig. E3. **Analysis of the small molecules partitioning within liquid droplets containing 45% w/v BSA. a**, Partitioning analysis of the small molecules pyruvate and NADH within the droplets phase and continuous phase (without adding enzymes during droplets preparation) using droplets with 45% BSA (w/v). The concentration of pyruvate and NADH in the two phases has been measured enzymatically by using LDH after separation of the two phases with a centrifugation step (30 minutes 16000 g). **b**, Partitioning analysis of ethanol and NADH within droplets and continuous phase using droplets with 45% BSA (w/v). The concentration of ethanol in the two separated phases has been measured enzymatically by using ADH while NADH concentration has been measured using LDH activity assay as reported in the Methods (LDH activity measurement).

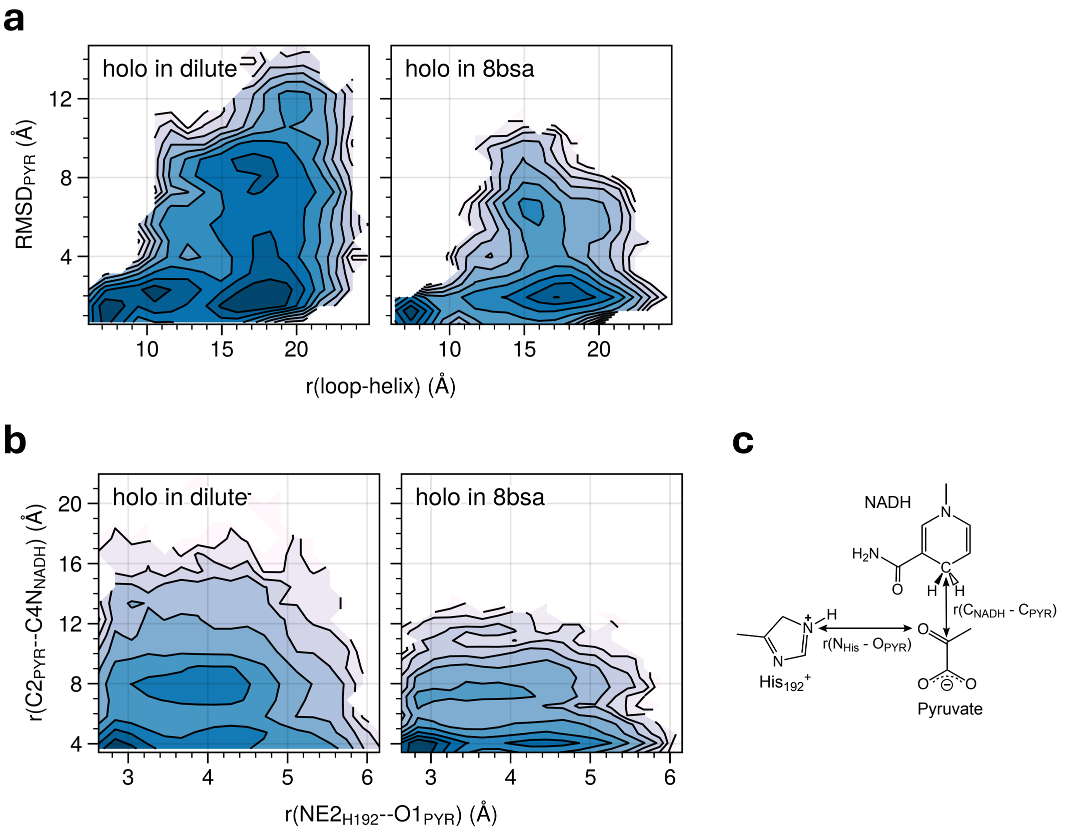


Extended Data Fig. E4. **Two dimensional PMF in dilute and crowded (8bsa) conditions.** Plots in terms of **a** active site loop and its contacting helix and RMSD of pyruvate, and **b** the distance between an oxygen atom of pyruvate and a nitrogen atom of His192 and the distance between a carbon atom of pyruvate and a carbon atom of NADH shown in **c**.

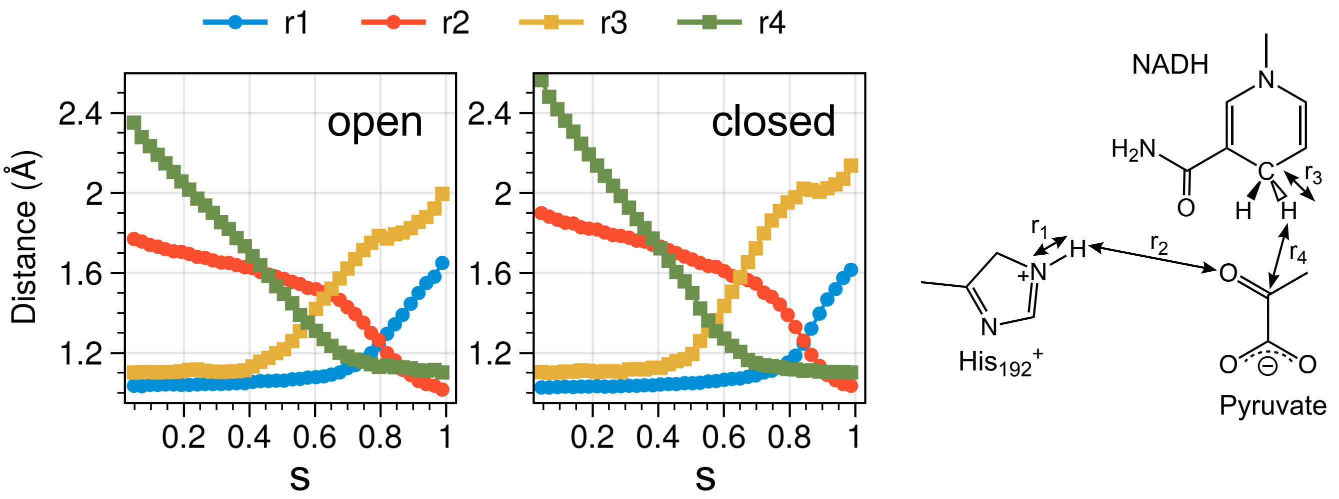


Extended Data Fig. E5. **The variation of bond length along the reaction path (s; normalized pathCV) in the open and closed forms of LDH monomers.** $r_1$ and $r_2$ are the breaking and forming bonds in the PT reaction, while $r_3$ and $r_4$ are those in the HT reaction.
